# Molecular basis for the R-type anion channel QUAC1 activity in guard cells

**DOI:** 10.1101/2021.09.09.459598

**Authors:** Li Qin, Ling-hui Tang, Jia-shu Xu, Xian-hui Zhang, Yun Zhu, Chun-rui Zhang, Xue-lei Liu, Mei-hua Wang, Fei Li, Fei Sun, Min Su, Yu-jia Zhai, Yu-hang Chen

## Abstract

The rapid (R)-type anion channel plays a central role in controlling stomatal closure in plant guard cells, thus regulating the exchange of water and photosynthetic gas (CO_2_) in response to environmental stimuli. The activity of the R- type anion channel is regulated by malate. However, the molecular basis of the R-type anion channel activity remains elusive. Here, we describe the first cryo-EM structure of the R-type anion channel QUAC1 at 3.5 Å resolution in the presence of malate. The structure reveals that the QUAC1 is a symmetrical dimer, forming a single electropositive T-shaped pore for passing anions across the membrane. The transmembrane and cytoplasmic domains are assembled into a twisted bi-layer architecture, with the associated dimeric interfaces nearly perpendicular. Our structural and functional analyses reveal that QUAC1 functions as an inward rectifying anion channel and suggests a mechanism for malate-mediated channel activation. Altogether, our study uncovers the molecular basis for a novel class of anion channels and provides insights into the gating and modulation of the R-type anion channel.

## INTRODUCTION

Stomatal pore, formed by a pair of guard cells in leaf epidermis, plays a crucial role in regulating CO_2_ assimilation and water evaporation in plants ^1,2^. Guard cells can integrate a wide range of environmental stimuli, such as drought, humidity, high CO_2_, and ozone, and convert them into appropriate turgor pressure changes that regulate stomatal opening or closure ^3,4^.

It is well-established that anion effluxes are key events to initiate stomatal closure in response to environmental stimuli ^5–8^. Previous studies showed that the guard cells harbour two distinct types of anion channels: rapid (R)-type and slow (S)-type ^9–12^. The R-type anion channel can be fully activated and rapidly inactivated within milliseconds, whereas the S-type anion channel takes up to several tens of seconds to be activated, followed by slow inactivation kinetics ^10^. Recent works of <u>SL</u>ow <u>A</u>nion <u>C</u>hannel 1 (SLAC1), an S-type anion channel discovered in *Arabidopsis* ^13–15^, have uncovered detailed mechanisms underlying S-type anion channel activity ^16–21^. However, the molecular basis of the biological role of the R-type anion channel is still largely enigmatic due to the lack of structural information.

<u>QU</u>ick <u>A</u>nion <u>C</u>hannel 1 (QUAC1), initially named aluminium-activated malate transporter12 (ALMT12), represents a guard cell R-type anion channel in *Arabidopsis* ^22^. The loss-of-function of *quac1* (*almt12*) mutation resulted in impaired stomatal closure ^22,23^. A double mutant lacking SLAC1 and QUAC1 was nearly unresponsive to various stimuli that were expected to cause stomatal closure ^24^. Apart from its role in stomatal regulation, QUAC1 also plays a vital role in cell polarity and the growth of pollen tubes via regulation of apical anion efflux ^25,26^. The activity of QUAC1 was reported to be modulated by malate ^22,27–29^, OST1 phosphorylation ^30^ and calmodulin ^31^, but the underlying mechanisms remain elusive.

Here, we report the first cryo-EM structure of soybean QUAC1 from Glycine max (*Gm*QUAC1, ~67% sequence identity to Arabidopsis *At*QUAC1) at the resolution of 3.5 Å in the presence of malate. The structure revealed that the QUAC1 is a symmetrical dimer assembled into twisted bi-layer architecture with a central T-shaped pore across the membrane. The pore is lined with highly conserved positively charged residues, rendering the pore surface electropositive, consistent with the QUAC1 function as an anion channel. Further electrophysiological analyses reveal that the QUAC1 displays rapid kinetics in channel gating during activation and inactivation.

Our truncation study reveals that the N-terminal juxtamembrane helix is crucial in maintaining structural integrity for proper channel function. Further mutagenesis analyses demonstrate that the C-terminal interaction locks the channel in a high-energy state, and that a domain-swapped helix plays an inhibitory role in channel regulation. Two pore-lining W90 residues, close to the lateral fenestration within the transmembrane domain, serve as toggle switches to regulate gating during channel activation. Therefore, the unique architecture endows QUAC1 function as an R-type anion channel, and our structural and functional analyses allow us to propose a novel mechanism for malate-mediated channel activation. Altogether, our study uncovers the molecular basis for a novel class of anion channels and provides insights into the gating and modulation of the R-type anion channel.

## RESULTS

### Bioinformatics analyses of plant ALMTs/QUACs

QUAC1 belongs to the ALMT family, which has diverse functions in plants, including stomatal function ^22,23,32–34^, pollen tube growth ^25,26^, Al^3+^ resistance ^35–38^, mineral nutrition ^39,40^, fruit acidity ^41,42^, microbe interactions ^43,44^, and seed development ^45,46^. There are thirteen members in the model plant *Arabidopsis thaliana*, except for one short sequence of a partial transmembrane domain (ALMT11). To better understand how QUAC (ALMT) proteins are represented in the plant kingdom, we clustered ~1,700 non-redundant sequences into a superfamily at the PSI-BLAST level E≤5×10^−3^, then into two distinct families at a threshold of E≤10^−160^, and finally into subfamilies at a threshold of E≤10^−180^, as detailed in Extended Data Table 1.

Since more studies have shown that ALMT members function as anion channels rather than transporters, and the QUAC1 is now the best-characterized member, we adopted nomenclature for the QUAC superfamily, which was further divided into families QF1 and QF2. We further found that family QF1 has two subfamilies: QF1A comprises the *Arabidopsis* ALMT3, 4, 5, 6, 9 and their homologs, and QF1B has a distinct set of ALMT proteins. Family QF2 has three subfamilies: the *Arabidopsis* ALMT1, 2, 7, 8, 10 proteins are in subfamily QF2A, the *Arabidopsis* ALMT12, 13, 14 proteins are in QF2B, and another distinct set of ALMT proteins are in QF2C (Extended Data Fig. 1).

The structure-based sequence alignment for the *Gm*QUAC1 and thirteen *Arabidopsis* QUAC/ALMTs is shown in Extended Data Fig. 2. Sequence analyses reveal that the ALMTs share relatively conserved transmembrane and cytosolic helical regions. Some conserved positively charged residues are distributed in the transmembrane regions and may be responsible for anion selectivity or voltage sensing. Some fingerprint motifs, including WEP (Trp-Glu-Pro) and PXWXG (Pro-X-Trp-X-Gly), are found in the cytosolic helical region. One of the most divergent regions (~50-100 aa in length) is found between H5 and H6 helices in the cytosolic portions, harbouring potential phosphorylation residues (Ser/Thr) for channel modulation. Further studies will provide mechanistic insight into the regulation of these characteristic motifs or distinct sets of QUACs.

### Cryo-EM structural determination of QUAC1

We screened six plant QUAC1s for expression and found that the *Gm*QUAC1 was suitable for further structural and functional studies (see method for details). Upon solubilization in 1.0% n-dodecyl-β-D-maltopyranoside (DDM) and 0.02% cholesteryl hemisuccinate (CHS), the *Gm*QUAC1 proteins were purified by Ni^2+^-affinity chromatography, and the resulting peak-fractions were pooled and loaded onto a gel-filtration column for further purification, buffer and detergent exchange. The final solution contained 0.005% detergent LMNG, with either 150 mM NaCl or 75 mM L-malate.

We further determined its cryo-EM structure in the presence of 75 mM L-malate (Extended Data Fig. 3, 4 and Extended Data Table 2). The dimeric reconstruction at 3.5 Å resolution allowed *de novo* modelling of 410 of the 537 amino acids (residues 36-393 and 455-506) per protomer chain. Other regions are not resolved in the density map due to their intrinsic flexibility in the protein. The search in the Dali server ^47^ for similar structures returned no significant hit. To the best of our knowledge, the QUAC1 protomer has a unique protein fold and architecture, representing a novel class of ion channels.

### The QUAC1 channel forms a twisted bi-layer architecture

The QUAC1 channel is a flat vase-shaped homodimer with a two-fold axis perpendicular to the plasma membrane (Fig. 1a and Supplementary Video 1). The overall molecule forms a bi-layer architecture, divided into two portions of the transmembrane domain (TMD) and the cytoplasmic helical domain (CHD). The domain topology for each protomer is shown in Fig. 1b. The TMD comprises 6 TM helices, arranging as three pairs of V-shaped helical hairpins stacked against each other, whereas the CHD has 7 helices, forming a helical bundle with a short domain-swapped finger helix. The N-terminal pre-TM region contains a juxtamembrane helix bent at a conserved proline before the TMD. A disorder region between helix H5 and H6 in the CHD enriches with Ser/Thr residues (Extended Data Fig. 2), raising an intriguing possibility of containing phosphorylation sites for channel regulation.

The dimer formation is mediated by interactions from both TMDs and CHDs, which bury surface areas of ~3,500 Å^2^ and ~3,300 Å^2^, respectively. When viewed from the extracellular side, the TM helices trace the circumference of an ellipse with TM1-TM6 from one protomer followed by another set of TM1-TM6 from the second protomer in an antiparallel manner (Fig. 1c). At the membrane, TM1 and TM2 directly interact with TM4 and TM5 from another protomer, respectively. In the cytoplasm, the CHD of each protomer unites together at the distal end of the molecule, mediated by intensive interactions. A domain-swapped finger helix (H6) also participates in C-terminal interactions (Fig. 1d). However, the functional role of these interactions remains unclear.

**Fig. 1.**
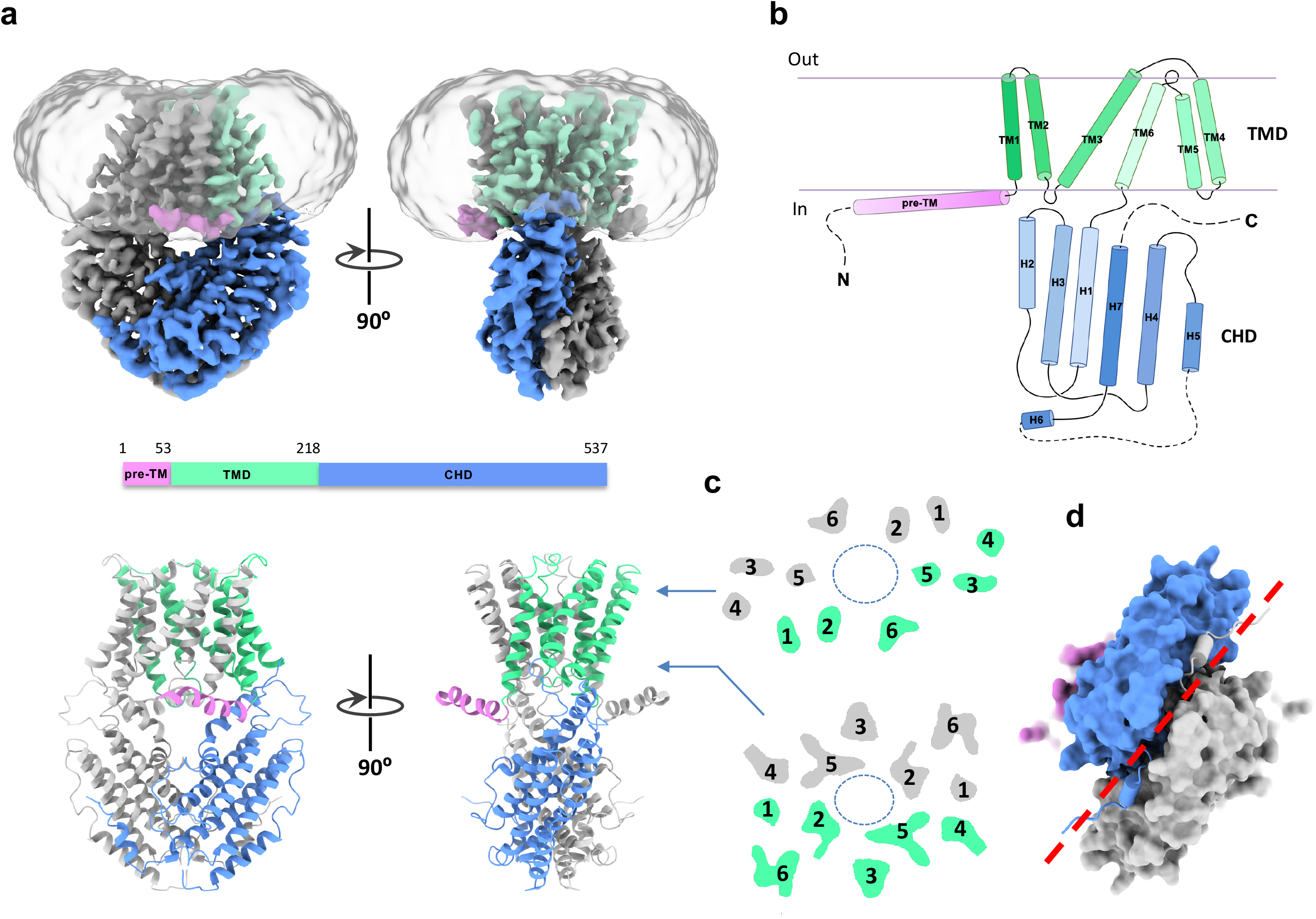
Architechture of the QUAC1 channel. **a**, Overall structure of the *Gm*QUAC1 channel. The cryo-EM density maps (top) and ribbons drawings (bottom) are shown, with the pre-TM helix in pink, the transmembrane domain (TMD) in green and the cytoplasmic helical domain (CHD) in blue in one promoter. Another promoter is coloured in grey. **b,** Topology of the *Gm*QUAC1 protomer. The six helices in TMD are marked by TM1-6, and the seven helices in CHD are marked as H1-7. The membrane boundary is shown as grey lines. The disordered regions are indicated as dashed lines. **c,** Top view of the cross-section of the transmembrane layer at the indicated positions by the arrow lines in (**a**). The elliptical dashed line marks the pore, and the TMs are indicated. **d,** Bottom view of the C-terminal dimeric domains, colored as in (**a**). The domain-swapped finger helix H6 is shown as cylinder cartoon, and a red dash line marks the dimeric interface of the CHD.

In the bi-layer structure of QUAC1, the TMDs connect to the CHDs via a highly conserved PXWXG motif (Pro-X-Trp-X-Gly). They interact with each other in a twisted manner, with their dimeric interfaces nearly perpendicular to each other (forming a dihedral angle of ~83°) (Fig. 2a). Interestingly, another characteristic motif WEP (Trp-Glu-Pro), located in the connecting loop between helix H2 and H3, is found in the vicinity of the PXWXG motif at the layer interface between TMD and CHD. These two motifs, together with other surrounding charged residues (Arg/Lys: R56, K63, R291, and R294; Asp/Glu: D54, E223, D224, and E289), form intensive interactions at the layer interface, and thus may play a vital role in coupling channel regulation to gating (Fig. 2b). Mutations in the WEP motif (E286Q in *Ta*ALMT1 and E276Q in *At*QUAC1) ^27,48^completely abolish their channel activities, possibly due to impaired coupling between the TMD and CHD caused by the altered interaction at the layer interface. Thus, the unique architecture design causes the channel in a high-energy state, providing a structural basis for QUAC1 function as an R-type channel. These observations also imply that the intramolecular domain rearrangements may occur upon channel activation.

**Fig. 2.**
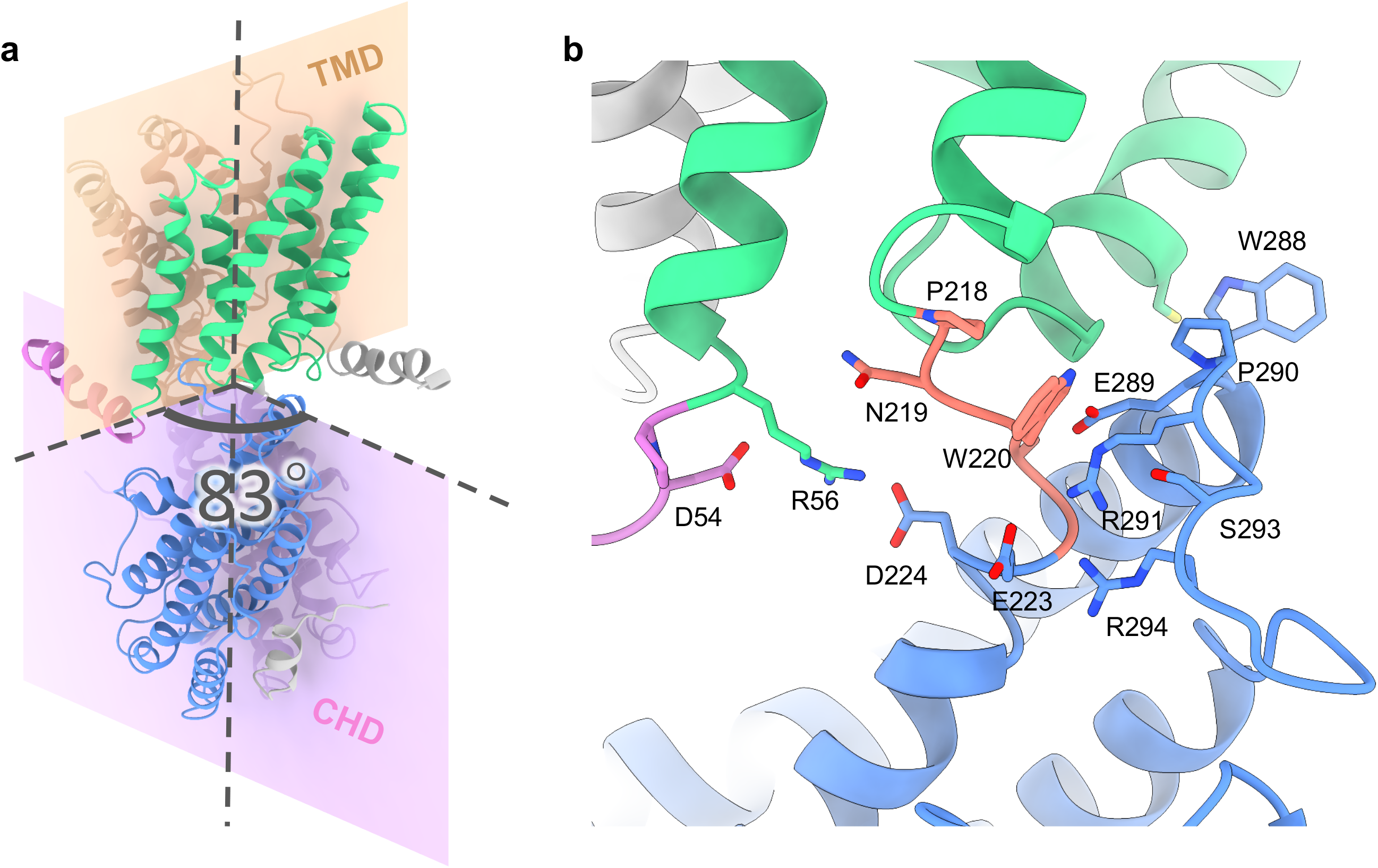
The twisted bi-layer architecture of the QUAC1 channel. **a,** The dimeric interfaces of the TMD and CHD of the *Gm*QUAC1 channel. These dimeric interfaces are nearly perpendicular to each other, forming a dihedral angle of ~83°. The ribbon model is colored as in Fig. **1a**. **b,** The layer interface between the TMD and CHD. The TMD portion connects to the CHD portion via a highly conserved PXWXG motif (P218-N219-W220-S221-G222, in salmon), which is interact with another characteristic motif WEP (W288-E289-P290, located between helix H2 and H3, in blue). Other surrounding charged residues at the interface are also shown in sticks, including R56, R291, R294, D54, E223, D224 and E289. The ribbon model is colored as in (**a**).

### Structural features of the ion conduction pathway in QUAC1

Unlike the trimeric SLAC1, which has three independent pores ^20^, the dimeric QUAC1 forms a single T-shaped tunnel with a bifurcated entrance in the cytoplasm (Fig. 3a). The TM2, TM3, TM5, and TM6 from both protomers create a pore with a radius of ~4-6 Å within the membrane (Fig. 3b), as estimated by HOLE ^49^. Overall, the QUAC1 pore is lined with highly conserved and general hydrophobic residues (Fig. 3c). Nevertheless, some highly conserved positively charged residues are distributed along the pore, thus rendering the surface electropositive (Fig. 3d).

**Fig. 3.**
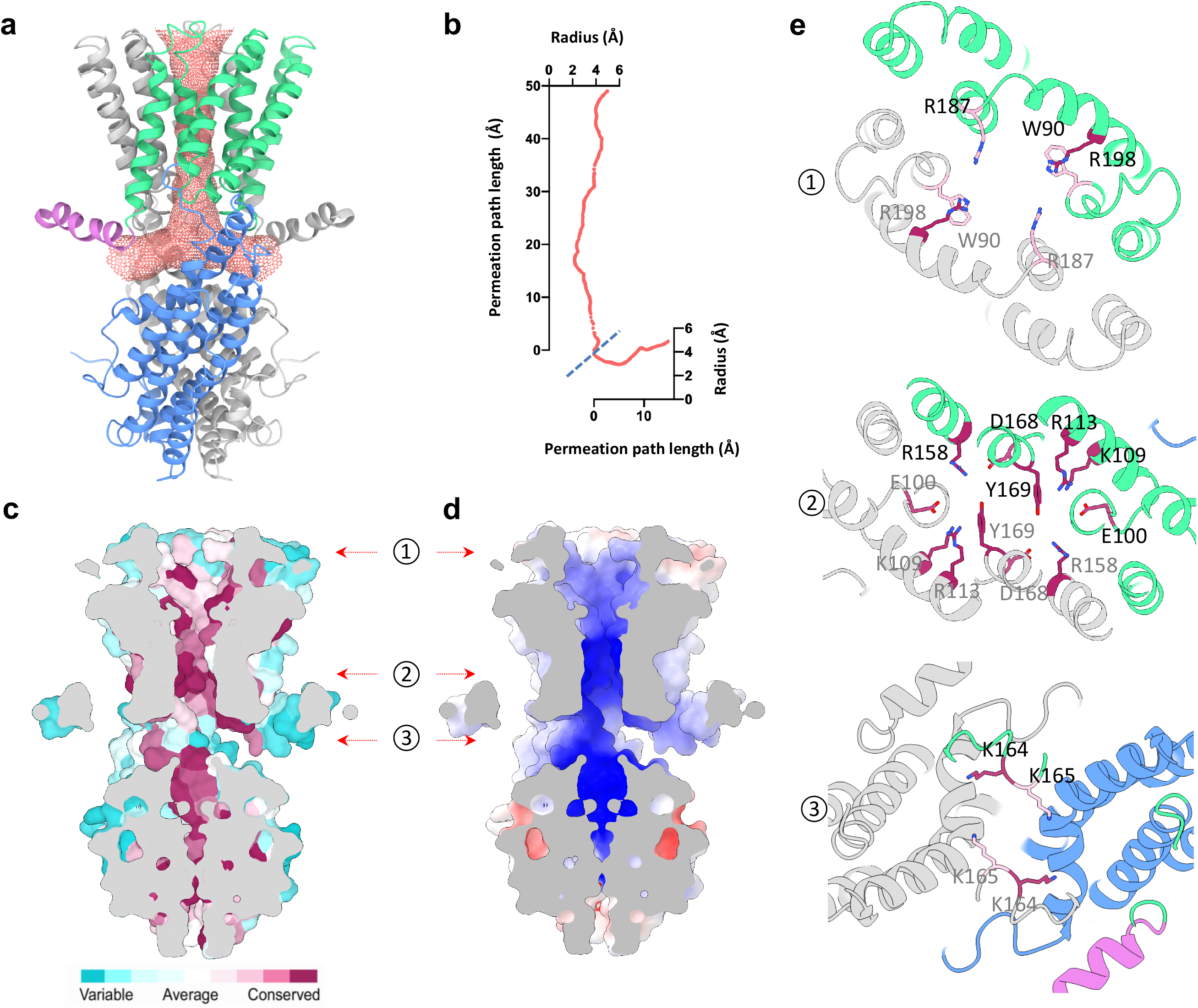
Structural feature of the QUAC1 channel pore. **a,** The pore-lining surface was computed by the program HOLE and drawn into a ribbon model of the *Gm*QUAC1 pore. We used a simple van der Waals surface for the protein and the program default probe radius of 1.15 Å. The T-shaped pore is shown in salmon dots. **b,** Plot of the pore radius as a function of the pore axis. **c** and **d,** Side views of the cross-section through the ion-conducting pore. The molecular surface colored with sequence conservation for 137 non-redundant proteins of the QF2B subfamily is shown in (**c**) and the molecular surface colored with electrostatic potential is shown in (**d**). **e,** Top views of the cross-section through the ion-conducting pore at three indicated positions by the arrow in (**c**) and (**d**). The ribbon is colored as in (**a**). The conserved pore-lining charged residues are shown in sticks, coloured with the sequence conservation as in (**c**).

Two arginines (R187 and R198) form a positively charged ring facing the pore entrance at the extracellular side. Three positively charged residues from TM3 and TM4 (K109, R113, and R158) protrude into the pore, making a second positively charged ring within the membrane. They are associated with other conserved residues from TM2 and TM5 (E100, D168, and Y169), mediating an inter-helices network within the pore. Another two lysines (K164 and K165) form additional constrictions at the bottom of the T-shaped pore (Fig. 3e). Altogether, the structural feature of QUAC1 likely contributes to its function as an anion-conducting channel. The above findings suggest that permeable anions interact with the pore-lining charged residues, thus participating in channel regulation.

Remarkably, a central kink (~24°) in TM6 causes a lateral fenestration from the membrane to the lumen of the channel pore within each protomer (Fig. 4a). The fenestration measures ~6 Å × 20 Å, and its dimension depends upon the conformation of pore-forming TMs, especially the kinked TM6. The presence of kinked helix and lateral fenestration in the structure implies that the channel in 75 mM malate is in a constrained conformation. Interestingly, several unmodeled densities and a highly conserved pore-lining W90 residue are found within or near the fenestration (Fig. 4b), raising an interesting question if the bulky W90 residue serves as a toggle switch to regulate channel gating during QUAC1 activation.

**Fig. 4.**
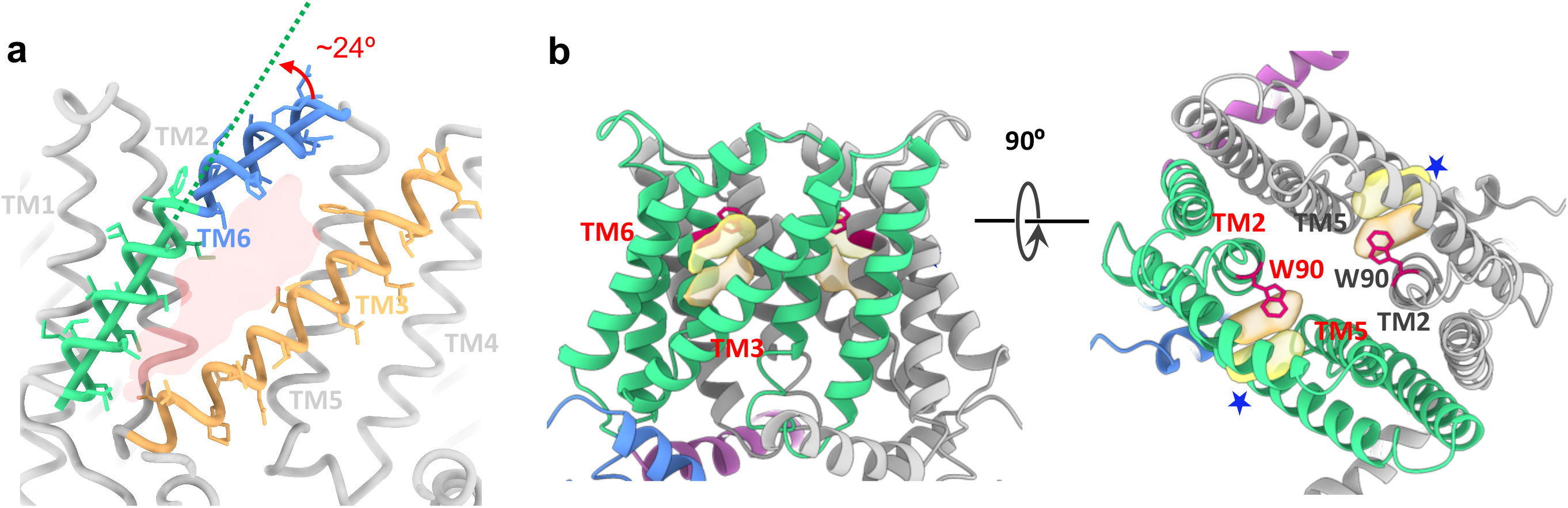
The lateral fenestration in the QUAC1 channel. **a,** The lateral fenestration, with a dimension of ~6 Å × 20 Å, is shown in transparent red, which is formed by TM2, TM3, TM5 and TM6 from one protomer. A kink (~24°) in TM6 is indicated. **b,** The unidentified densities (in yellow) within the fenestration (indicated by arrows). The side view (right) and top view (left) are shown. The W90 residues are shown as red sticks. The locations of fenestration are indicated with a bule stars in the top view.

### Functional characterization of QUAC1 by TEVC

A total of 34 ALMTs were found in soybean ^50^, but none of them has yet been functionally characterized by electrophysiology. We expressed the *Gm*QUAC1/ALMT12 in *Xenopus laevis* oocytes and measured channel conductance by applying the two-electrode voltage-clamp (TEVC) technique. In the recordings, the application of voltage pulses elicited rapidly activating currents in the bath solution of malate, resembling the *At*QUAC1 ^22^. We also found that the steady-state currents mediated by *Gm*QUAC1 displayed a bell-shaped current-voltage curve (Extended Data Fig. 5). This electrophysiological behaviour points to a strong voltage dependence for the QUAC1 channel activation, a hallmark feature of the R-type anion channel in guard cells ^10,22^.

We also measured the conductance of *Gm*QUAC1 in other bath solutions. Surprisingly, we found that the activation/inactivation kinetics and voltage-dependency of *Gm*QUAC1 in NaNO_3_ or NaCl solutions are distinguished from those in malate solution (Extended Data Fig. 5). In contrast to substantial instantaneous currents, only small steady-state currents can be seen in the recording of *Gm*QUAC1 in the bath solution of NaNO_3_ or NaCl (Extended Data Fig. 5). Overall, our TEVC recordings demonstrate that external anions play a crucial role in regulating QUAC1 channel properties, especially the activation/inactivation kinetics and steady-state currents. We hypothesize that the anions interplay with the pore-lining residues, such as the inter-helices charged residues interacting network within the pore (Fig. 3e), and act as door stopper wedges to regulate channel gating.

### Single-channel analysis of QUAC1 channel in planar lipid bilayer

To further understand the single-channel properties, we first fused the *Gm*QUAC1^NaCl^ proteins (purified in 150 mM NaCl) into a lipid bilayer and measured channel conductance under different symmetrical solutions (Fig. 5a). When using symmetrical NaNO_3_ solutions, the application of a series of transmembrane voltages to the *trans* chamber resulted in frequent openings and closings of the *Gm*QUAC1 channel, as evidenced by the current fluctuation in the recording traces at different applied voltages (Extended Data Fig. 6a). We also found that the channel gating transits between two predominant full open and closed states, and sometimes to a sub-conductance that is ~50% level of the full conductance (Fig. 5b). This observation is consistent with the two-entrance T-shaped tunnel presented in the QUAC1 channel (Fig. 3a). At ~50% of the full level, the intermediate sub-conductance suggested one of two entrances was blocked or closed. In addition, the QUAC1 channel gating displays a flickering feature that represents the rapid character of the activation/inaction kinetics for R-type anion channels.

**Fig. 5.**
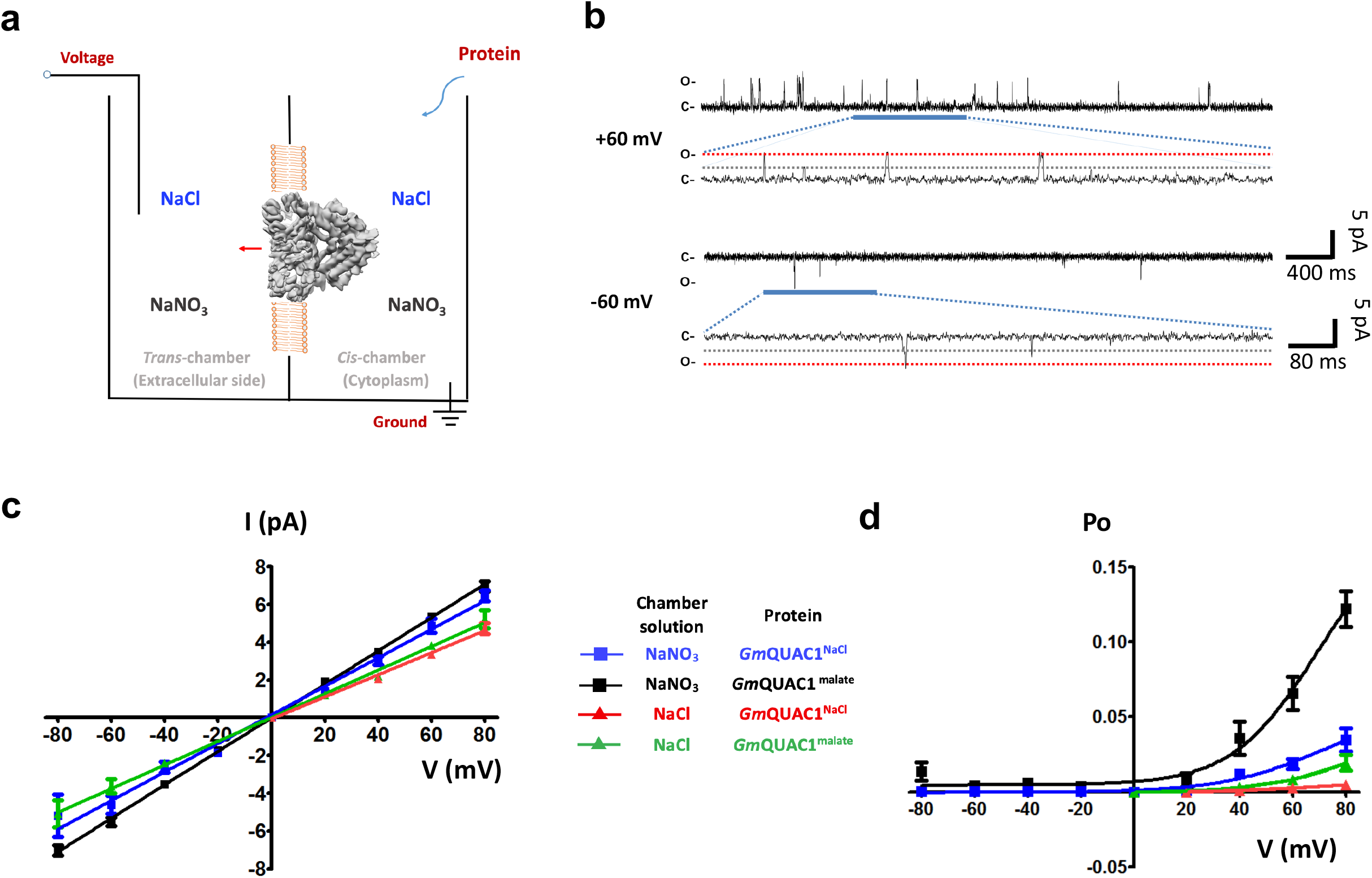
Single channel analysis of the QUAC1 channel in planar lipid bilayer. **a,** The schematic illustration of single-channel recording of *Gm*QUAC1 channel in a planar lipid bilayer. The chambers were filled with 1.0 ml of symmetrical solutions of 150 mM NaCl or 150 mM NaNO_3_, and the *Gm*QUAC1^NaCl^ or *Gm*QUAC1^malate^ (purified in 150 mM NaCl or 75 mM L-malate, respectively) were added to the *cis-*side. The *trans*-chamber, representing the extracellular compartment, was connected to the head stage input of a bilayer voltage-clamp amplifier. The *cis-*chamber, representing the cytoplasmic compartment, was held at virtual ground. **b,** Representative current traces for single channel analysis of *Gm*QUAC1^NaCl^ are shown at +60 mV or −60 mV holding potentials in the NaNO_3_ solution. The closed (C) and full-open (O) states are indicated. **c** and **d,** The current-voltage relationships (**c**) and open probabilities (**d**) for the single-channel recordings of *Gm*QUAC1. The chamber solutions (NaNO_3_ or NaCl) and protein (*Gm*QUAC1^NaCl^ or *Gm*QUAC1^malate^) are indicated in the inset (Data are mean ± SEM, n≥4).

In contrast, we observed much reduced channel activity in the measurements in the solution of 150 mM NaCl (Extended Data Fig. 6b). The mean single-channel conductance for the *Gm*QUAC1^NaCl^ is 75.5 ± 4.5 pS (n=5) and 58.5 ± 7.0 pS (n=5) in the solutions of NaNO_3_ and NaCl, respectively (Fig. 5c). Intriguingly, the channel open probability displays a strong voltage dependency, much higher at positive voltage than at negative one (Fig. 5d). As in our experimental setup, positive currents mean anions flow from the *cis* (representing cytoplasm) to the *trans*-chamber (representing extracellular side), and *vice versa*. Therefore, these results demonstrate that the QUAC1 channel functions as an R-type anion channel, favouring efflux (corresponding to anions flow from *cis-* to *trans-*chamber).

### Malate regulates QUAC1 channel gating

To investigate the effect of malate on the channel activity, we reproduced the same experiments with the *Gm*QUAC1^malate^ (purified in 75 mM L-malate) as above. Interestingly, we observed that *Gm*QUAC1^malate^ exhibits a similar current amplitude but a much higher open probability than those with *Gm*QUAC1^NaCl^ in both solutions of NaNO_3_ and NaCl (Fig. 5c and Extended Data Fig. 6). The mean single-channel conductance for the *Gm*QUAC1^malate^ is 88.5 ± 2.4 pS (n=4) and 62.7 ± 4.1 pS (n=4) in the solutions of NaNO_3_ and NaCl, respectively. The open probabilities of *Gm*QUAC1^malate^ also display a similar voltage dependency as observed for *Gm*QUAC1^NaCl^ (Figure 5D and S6). Taken together, our results suggest that malate regulates QUAC1 activation by modulating channel opening probability.

To further examine malate regulation on the channel gating, 2 mM L-malate was added to the solutions upon the *Gm*QUAC1^NaCl^ fused into a lipid bilayer. Interestingly, the channels became more active after adding 2 mM L-malate to either the *trans*- or *cis*-chamber (Fig. 6a and 6b). The current amplitude slightly increases, whereas the open probability becomes much higher (Fig. 6c and 6d). These observations confirm that malate can stimulate channel activity via enhanced open probability. Our findings also suggest that the malate regulation site may be located within the pore and can be accessed from either the extracellular or intracellular side of the membrane. We speculate that the *Gm*QUAC1 structure in the presence of 75 mM L-malate may represent an open state, although the malate-binding site remained unclear due to insufficient information.

**Fig. 6.**
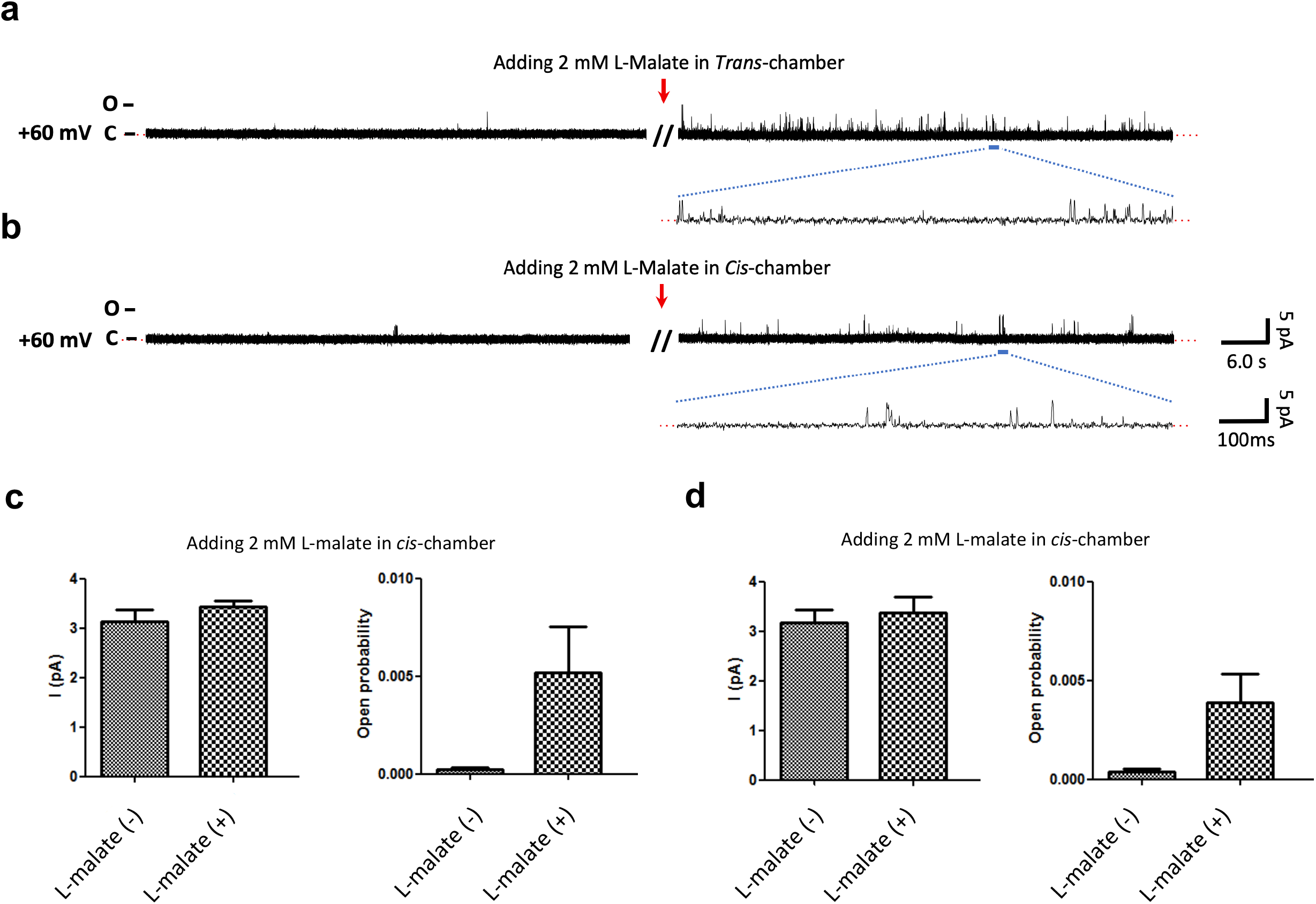
Single channel analysis of malate regulation on QUAC1. **a** and **b,** The chambers were filled with symmetrical solutions of 150 mM NaCl, and the purified *Gm*QUAC1^NaCl^ proteins were added to the *cis-*side. The representative current traces of 60 s at +60 mV, before and after adding 2 mM L-malate to the *trans*-(**a**) or *cis*-(**b**) chamber, are shown. **c** and **d,** The regulation effect of malate on the *Gm*QUAC1^NaCl^ in the NaCl solutions. The current amplitude (left) and open probability (right) analysis for the recordings, before (−) or after (+) addition of 2 mM L-malate to the *trans*-(**c**) or *cis*-(**d**) chamber (Data are mean ± SEM, n=3 for each group).

### The N-terminal pre-TM helix is required for channel activity

The sequence analysis shows that the N-terminal segment before TMD is one of the most divergent regions in the QUAC family (Extended Data Fig. 2). The QUAC1 structure reveals that this region contains a juxtamembrane pre-TM helix (Fig. 1b), enriching positively charged residues. To examine its functional role, we designed a truncation construct (Δ1-53). Surprisingly, we found that the Δ1-53 mutant resulted in nearly null currents in the TEVC recordings (Extended Data Fig. 7). The observation demonstrates that the pre-TM helix is indispensable for channel activity. Considering its association with the inner leaflet of the membrane, we propose that the pre-TM helix may serve as a lever to regulate the proper conformation of the QUAC1 channel.

### The C-terminal dimeric interaction is crucial for channel function

The cytosolic C-terminal portion is mainly formed by α-helices, except that a large divergent region between helix H5 and H6 is disordered or missing in the QUAC family (Extended Data Fig. 2). The *Gm*QUAC1 structure revealed extensive interactions between two CHDs at the distal end of the dimer (Fig. 7a). Two highly conserved hydrophobic residues (F470 and L474 from helix H7) protrude into the hydrophobic pocket formed by another protomer. Together with their antiparallel counterparts from another protomer, these bulky hydrophobic residues form a zipper-like interaction at the dimeric interface (Fig. 7b). In addition, a short finger helix (H6), whose sequence is conserved in the subfamily QF2B (Extended Data Fig. 2), also participates in the dimeric interaction in a domain-swapped manner (Fig. 7c). These interactions unite the two CHDs together and thus creates an anchor point for mechanical force transduction between TMD and CHD.

**Fig. 7.**
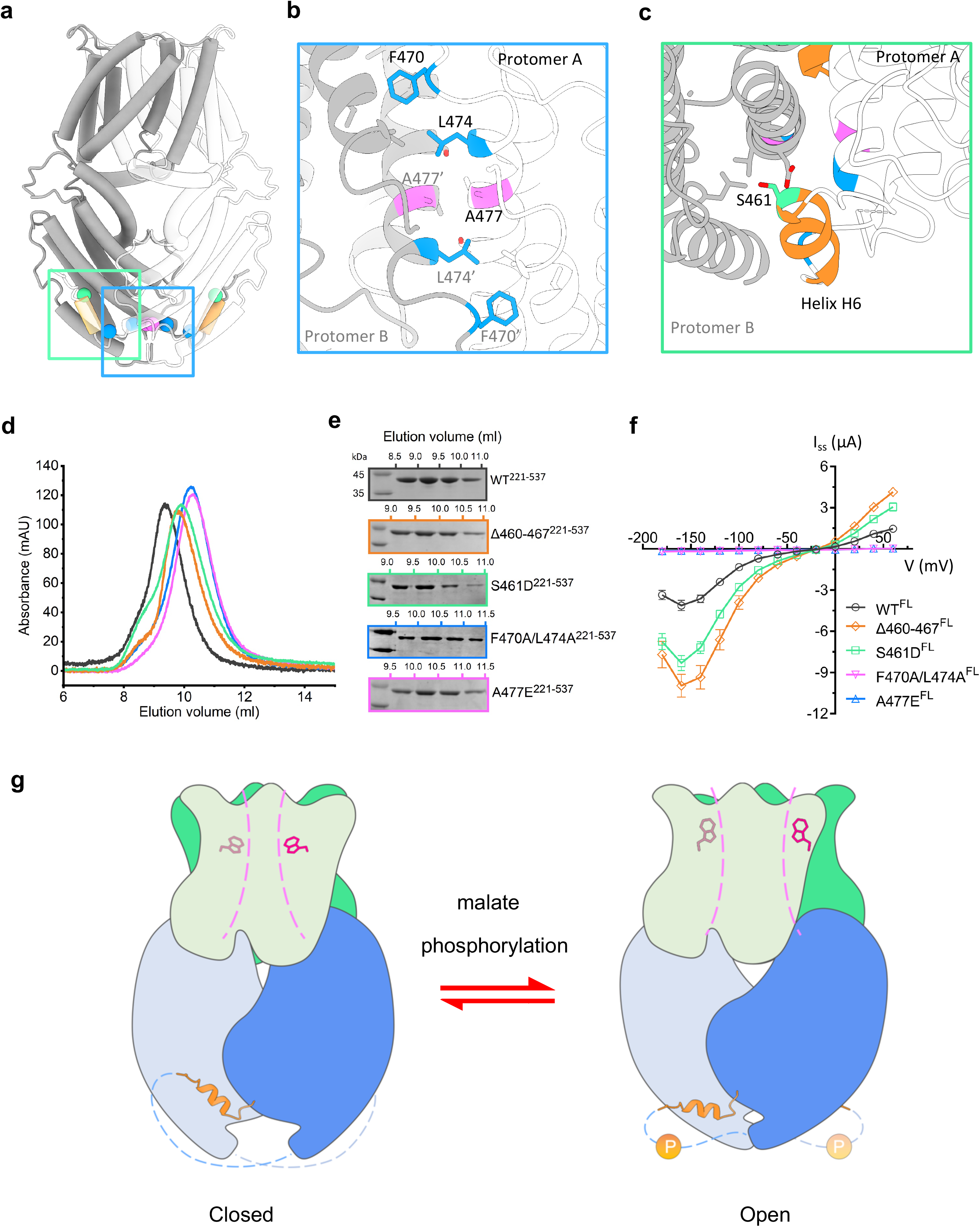
Effect of dimeric interactions on channel activity and the proposed model for QUAC1 regulation. **a,** The cartoon of the *Gm*QUAC1 channel is shown, with one protomer in grey and one in white. The domain-swapped helix (H6, residues 460-467) is colored in orange, and key residues at the dimeric interface are highlighted: S461 in green, A477 in magenta and F470/L474 in blue. **b** and **c,** Close-up views of the cytoplasmic dimer interface. Residues at the dimeric interface are shown as sticks and colored as in (**a**). **d** and **e,** The purification of the CHD proteins (residues 221-537, wild-type or mutants) on a Superdex75 (10/300) column. Compared to the wild-type (peak at ~9.2 ml), the eluted peaks for the dimer-disrupting mutants (A477E and F470A/L474A) shift backwards to ~10.2 ml, suggesting a disassociation of dimer into monomer. The elution peaks at ~9.7 ml for the two finger helix altered mutants (Δ460-467 and S461D) suggest loosened conformations upon the removal of finger helix interaction. The eluted fractions were also analyzed by SDS-PAGE, as shown in (**e**). **f,** The TEVC recording of *Gm*QUAC1 channel (wild-type and mutants) in *Xenopus laevis* oocytes. The same set of mutations, as in (**d**), were generated into full-length constructs for conductance measurement in the bath solution of 30 mM L-malate (Data are mean ± SEM, n≥8). **g,** Model for the QUAC1 channel regulation. Upon malate binding, the QUAC1 undergoes intramolecular domain reorganization to form a twisted bi-layer architecture, and subsequently induces the conformational change of the toggle switch W90. These changes promote the channel conversion from closed to open state. Other modulations, such as phosphorylation or calmodulin binding, can further enhance the malate-mediated activation, probably via the release of its inhibitory domain-swapped helix from the inter-protomer interaction.

To investigate the functional role of the dimeric interactions in channel regulation, we generated mutations, aiming to disrupt the dimer-interface (A477E and F470A/L474A), or to break the finger helix interaction in H6 (Δ460-467 and S461D), in both the CHD (residues 221-537) bacteria-expression constructs for gel-filtration analysis and the full-length oocyte-expression constructs for conductance measurement. In contrast to the wild type, both dimer-interface disrupting mutants exhibited backwards-shifted elution peaks (from 9.2 ml to 10.2 ml) in the gel filtration profile and showed almost null currents in TEVC recording. In another experiment, the two altered helix H6 mutants cause loosened dimers but unexpectedly enhanced channel activities (Fig. 7d-f and Extended Data Fig. 8). Altogether, our results show that the C-terminal interaction has two important functions: (1) joins the two CHD together and locks the channel in a high-energy state; (2) a domain-swapped helix (H6) plays an inhibitory role, and its release enhances the channel activity. Our results demonstrated that the C-terminal interaction provides a basis for fast regulation of the R-type anion channel.

### Gating of the QUAC1 channel

Our electrophysiological experiments have demonstrated that malate plays a vital role in modulating *Gm*QUAC1 activity; however, we wonder how malate regulation is coupled to channel activation. The *Gm*QUAC1 structure reveals that the conserved pore-lining tryptophan W90 is in close contact with an unmodelled density within the fenestration (Fig. 4b). To understand the functional role of W90, we made mutations of W90A and W90F. Compared to wild-type, the removal of indole ring in W90A mutant causes ~5 times larger conductance in the presence of external malate. A similar aromatic amino acid substitution in the W90F mutant can stimulate channel currents even larger (Extended Data Fig. 9a and 9b). These observations imply that the conserved W90 may interact with surrounding residues and function as a toggle switch to regulate channel gating.

This notion is confirmed in the measurement of combined mutants of W90F with other two loss-of-function mutants, Δ1-53 and A477E. We observed that both null mutants restore channel activity to a moderate level after the addition of W90F (Extended Data Fig. 9c and 9d). We speculate that malate promotes QUAC1 activation probably via regulating the conformation of tryptophan W90. Distinct from the S-type SLAC1 regulation controlled by phosphorylation/ dephosphorylation ^16–20^, our study reveals that the R-type QUAC1 gating can be directly mediated by malate, which is much faster. These findings may explain the fast kinetics observed in the R-type channel, compared with the slow activity found in the S-type channel.

## DISCUSSION

Recently, AlphaFold has made significant progress in 3D structure prediction from protein sequence ^51^. By taking advantage of this advanced technology, we generated an AlphaFold predicted model for the *Gm*QUAC1 protein that is the focus of our current study. Surprisingly, the AlphaFold predicted *Gm*QUAC1 structure (AF-structure) is surprisingly similar to that determined by cryo-EM (cryo-EM structure), with an r.m.s.d. of 2.1 Å/160 C_α_ and 1.4 Å/207 C_α_ for the TMD and CHD, respectively. However, further superimposition reveals a noticeable difference in the orientation of TMD vs CHD in QUAC1, with an r.m.s.d. of 3.1 Å/388 C_α_ and 3.2 Å/776 C_α_ for the monomer and dimer, respectively (Extended Data Fig. 10a).

Unlike the kinked TM6 observed in the cryo-EM structure, the TM6 in the AF-structure adopts a straight conformation, eliminating the lateral fenestration in TMD (Extended Data Fig. 10b). As the malate regulation effect on the channel is not taken into account, the resulting predicted model may represent an apo-structure in a low energy state. The cryo-EM structure determined in the presence of malate reveals a high-energy twisted conformation, thus representing a malate-activated structure. These two structures allow us to make comparisons and gain insight into the gating mechanism of the QUAC1 channel. It seems that the pore-forming helices move slightly outward, as suggested by the porcupine plot on the Cα comparison between the AF-structure and the cryo-EM structure (Extended Data Fig. 10c). Another noticeable difference is the dihedral angle between the TMD and CHD interfaces, which is 86.5° and 83.0° for the AF-structure and cryo-EM structure, respectively (Extended Data Fig. 10d). This observation shows a slight rotation (~3.5°) between these two states, suggesting that domain reorganization occurs during the conformational conversion. We speculate that the domain reorganization is associated with malate regulation, and subsequently triggers conformational changes of the toggle switch in the pore to activate the channel (Extended Data Fig. 10e and and Supplementary Video 1).

Our work provides the first glimpse of the molecular structures of the R-type anion channel. Our study showed that the TMD and CHD portions in QUAC1 interact in a twisted manner, rendering the channel in a high-energy state. Based on our data, we propose the following mechanism for QUAC1 activation: upon malate binding, the QUAC1 undergoes intramolecular domain reorganization to form a twisted bi-layer architecture, which subsequently induces the conformational change of the toggle switch W90. These changes promote the channel conversion from the basal to the activated state. The malate-mediated activation is further enhanced by other modulations, such as kinase phosphorylation or calmodulin-binding, probably via releasing its inhibitory domain-swapped helix from the inter-protomer interaction (Fig. 7g and Supplementary Video 2). Our study provides mechanistic insights into the gating activation of the R-type channel and offers a plausible explanation for fast kinetics in the R-type currents.

As both SLAC1 and QUAC1 represent two essential distinct channels that mediate osmotic active anions efflux from guard cell, one remaining puzzle is how they interplay with each other to initiate stomatal closure in response to various environmental stimuli. Further investigation to address this issue will shed light on the understanding of guard cell stomatal signaling and provide critical information for engineering drought-resistant or water-use efficiency crops.

## SUPPLEMENTAL INFORMATION

Supplemental information can be found online at XXX

## ACKNOWLEDGMENTS

We thank the staff at the Center for Biological Imaging (CBI), Institute of Biophysics, Chinese Academy of Sciences. This project is financially supported by the National Key Research and Development Program of China (2020YFA0509903 and 2016YFA0500503 to Y.-h.C., and 2017YFA0504703 to Y.-j.Z.), the Strategic Priority Research Program of the Chinese Academy of Sciences (XDA24020305 to Y.-h.C., and XDB37040102 to F.S.), and the National Natural Science Foundation of China (31872721 to Y.-h.C., and 31771566 to Y.-j.Z.).

## AUTHOR CONTRIBUTIONS

L.Q. performed protein purification, Cryo-EM data collection, TEVC experiments, and data analysis; L.-h.T., J.-s.X., Y.Z. performed Cryo-EM data collection and structural determination; X.-h.Z. performed mutagenesis and protein purification; M.S. performed PLB experiments and data analysis; C.-r.Z., X.-l.L., M.-h.W, performed experiments; F.L., F.S., Y.-j.Z. analyzed data; Y.-h.C. initiated the project, planned and analyzed experiments, supervised the research and wrote the manuscript with input from all authors.

## METHODS

### Bioinformatics analysis of ALMT proteins

Sequences related to QUAC1/ALMTs were searched and analyzed by using PSI-BLAST ^52^. Searches at E<5*10^−3^ starting with thirteen individual *Arabidopsis* ALMTs identified a common pool of over 1,700 plant ALMT-related protein sequences, which were pooled together and used for sub-classification into families and subfamilies. Detailed information is reported in the footnotes to the Extended Data Table 1.

### Cloning and expression of QUAC1/ALMT12 in yeast *S. pombe*

Full-length coding sequences of six plant QUAC1s from rice, wheat, corn, soybean, tomato and cotton, were cloned into a modified pREP1 vector with a C-terminal 10× His tag. The resulting constructs were transformed into a leucine auxotrophic *S. pombe* strain, and the transformants were selected on the standard EMM plates without leucine, as previously described ^53^. To prevent protein expression during strain growth, 25 μM thiamine was added to inhibit the promoter.

### Scaled-up production and purification of *Gm*QUAC1

To prepare the seed, transformed cells were inoculated into 100 ml EMM culture medium supplemented with 25 μM thiamine and were shaken at 200 rpm and 30°C for 24 hr. For protein expression and scaled-up production, the seed cells were collected by centrifugation and washed with sterile water twice before inoculation to a culture of 500 ml. After 12 hr growth, 500 ml of fresh medium was supplemented, and the culture continued to grow for an additional 24 hr. Cells were harvested through centrifugation for 20 min at 4,500 rpm.

For protein purification, cells were re-suspended in lysis buffer (50 mM Tris-HCl pH 8.0, 200 mM NaCl, 2.5% glycerol, 1 μg/ml aprotinin, 1 μg/ml leupeptin, 1 μg/ml pepstatin, 2 mM PMSF and 2 mM DTT) and lysed using a high-pressure cell disrupter (JNBIO) with 4 passes at ~18,000 psi. Cell debris was removed by centrifugation at 12,000 rpm for 15 min, and the supernatant was subjected to a further ultra-centrifugation at 41,000 rpm for 1 hr. The membrane was collected and homogenized in a solubilization buffer (50 mM Tris-HCl pH 8.0, 200 mM NaCl and 2.5% glycerol) and incubated with a final concentration of 1.0% DDM and 0.02% CHS by gentle stirring for 1 hr at 4°C. After ultra-centrifugation at 35,000 rpm for 50 min, the resulting supernatant was purified by Ni^2+^-affinity column pre-equilibrated with the same solubilization buffer supplemented with 0.05% DDM and 2 mM TCEP. After 20 columns volume of buffer wash, the protein was eluted with 350 mM imidazole in the solubilization buffer. The resulting *Gm*QUAC1 protein, without removal of His-tag, was concentrated to ~10 mg/ml and loaded onto a Superose6 (10/300) gel-filtration column for further purification and detergent exchange. The gel-filtration buffer contained 20 mM Tris, 150 mM NaCl (pH 8.0, adjusted with HCl) or 75 mM L-malate (pH 8.0, adjusted with NaOH), and 0.005% LMNG. The elution fractions were analyzed by SDS-PAGE, and the peak fractions were collected and concentrated for functional analysis (~2 mg/ml) or making cryo-EM grids (~5 mg/ml).

### Cryo-EM grid preparation and data acquisition

4 μl of *Gm*QUAC1^malate^ proteins (purified in 75 mM L-malate) were applied to newly glow-discharged holy carbon film grids (Au R1.2/1.3, 300 meshes, Quantifoil, Germany). The grids were blotted with force 3 and blotting time of 9.0 s at 100% humidity and 4°C and were vitrified by plunge freezing into liquid ethane using Vitrobot Mark IV (Thermo Fisher Scientific, USA). All movies were collected on a Titan Krios G2 transmission electron microscope (Thermo Fisher Scientific, USA) operated at 300 KV, equipped with a Gatan K2 Summit direct detection camera (Gatan Company, USA) and a post-column GIF energy filter.

Data collection was performed on EF-TEM mode with a silt width of 20 e^−^·V. The magnification was set to a nominal 130,000x, corresponding to a calibrated pixel size of 1.04 Å/pixel at the specimen level (0.52 Å/pixel in super-resolution mode). Images were recorded using SerialEM (version 3.8.4) ^54^ with a beam-image shift method ^55^. During the 8s exposure, 32 frames were collected with a dose of around 60 e^−^/Å^2^. A total of 6,189 movie stacks were collected, with defocus ranging from −1.2 μm to −2.2 μm.

### Image processing and model building

For cryo-EM image processing, all main steps were performed using RELION 3.0 ^56^ or cryoSPARC 3.1 ^57^. Pyem ^58^ and UCSF Chimera ^59^ were used for format conversion of data files and reconstructions analysis, respectively.

In brief, 6,189 movie stacks were aligned by 5 × 5 patches with dose weighting and binned to 1.04 Å using Motioncor2 ^60^. The contrast transfer function parameters were estimated using GCTF ^61^. Micrographs with defects in the thon rings were discarded. 2,987,283 candidate particles were initially picked using Gautomatch without template ^62^, and followed by several rounds of 2D classification in cryoSPARC, giving a set of 2,136,754 high-quality particles. A rough initial model was generated by a subset of 100,000 particles. Subsequentially, six rounds of 3D classification were performed at C1 symmetry. Three parallel 3D classifications were performed within each round, and the resulting good particles were combined for sequential round classification. After that, a set of 428,433 particles that generated a map of 4.0 Å using NU-Refinement in cryoSPARC were subjected to Bayesian polishing. Three rounds of 3D classification were further performed as described below. Each round contains an NU-Refinement at C2 symmetry in cryoSPARC followed by an alignment-free 3D classification with a micelle-free mask in Relion. Finally, NU-refinement of 169,576 particles yielded a reconstruction with a resolution of 3.5 Å, based on the gold-standard Fourier Shell Correlation (FSC) using the 0.143 criterion. Density modification was performed using PHENIX ^63^. Local resolution was determined using ResMap ^64^.

A *de novo* atomic model was built manually in Coot ^65^ using the predicted secondary structure from MPI quick2d ^66^. The model was further refined using PHENIX ^63^. The final model was validated using EMRinger ^67^. All structure figures were prepared in UCSF ChimeraX ^68^.

### The two-electrode voltage-clamp recordings

The TEVC recordings were conducted as previously described ^20^. In brief, all cDNAs for *Gm*QUAC1, including wild-type or mutants, were cloned into plasmid pGHME2 for expression in *Xenopus laevis* oocyte. Linearised plasmids were used to generate cRNAs using T7 polymerase. 36 nanograms of cRNA of each construct were injected into isolated oocytes. Oocytes were then incubated at 18°C for ~48 hr in ND96 buffer (96 mM NaCl, 1.8 mM CaCl_2_, 1 mM MgCl_2_, 2 mM KCl, 5 mM HEPES-Na, pH 7.5).

Using a Oocyte Clamp OC-725C amplifier (Warner Instruments) and a Digidata 1550 B low-noise data acquisition system with pClamp software (Molecular Devices), TEVC recordings were performed in the bath solutions: 10 mM MES/Tris pH 5.6, 1mM Ca(gluconate)_2_, 1 mM Mg(gluconate)_2_, 70 mM Na(gluconate), 1 mM LaCl_3_, 30 mM NaCl or NaNO_3_, or 30 mM L-malate (pH 5.6, prepared from L-malic acid with NaOH). Osmolarity was adjusted to ~220 mOsmol*kg^−1^ with D-Sorbitol. The microelectrode solutions contained 3 M KCl, and the bath electrode was a 3 M KCl agar bridge. Voltage-clamp currents were measured in response to 200 ms-long voltage steps to test potentials that ranged from +60 mV to −180 mV in 20 mV decrement. Prior to each voltage step, the membrane was held at +60 mV for 50 ms, and following each voltage step, the membrane was returned to −180 mV for 50 ms. I-V relations for *Gm*QUAC1 channels were generated from currents measured at 5 ms after the beginning (Instantaneous currents, I_inst_) or before the end (Steady-state currents, I_ss_) of each test voltage step. Three independent batches of oocytes were investigated and showed consistent findings. Data from one representative oocyte batch were shown. The recordings were analyzed using Clampfit 10.6 (Molecular Devices) and Prism (ver. 5.0, GraphPad).

### The planar lipid bilayer recordings

Lipid bilayer experiments were conducted as previously described in our study of the ZAR1 channel ^69^. The purified *Gm*QUAC1 (in 150 mM NaCl or 75 mM L-malate), at a concentration of 1.5~2.5 μg/ml, was fused into planar lipid bilayers formed by painting a lipid mixture of phosphatidylethanolamine (DOPE) and phosphatidylcholine (DOPC) (Avanti Polar Lipids) in a 5:3 ratio in decane across a 200 μm hole in a polystyrene partition separating the internal and external solutions in polysulfonate cups (Warner Instruments). The *trans* chamber (1.0 ml), representing the extracellular compartment, was connected to the head stage input of a bilayer voltage-clamp amplifier. The *cis* chamber (1.0 ml), representing the cytoplasmic compartment, was held at virtual ground. Solutions used for I-V relationship analysis were as follows: 150 mM NaCl or 150 mM NaNO_3_. All solutions were buffered with 20 mM Tris (pH 8.0). Purified proteins were added to the *cis* side and fused with the lipid bilayer.

Currents were recorded every 1~2 min after application of the voltage to the *trans* side. Single-channel currents were recorded using a Bilayer Clamp BC-525D (Warner Instruments), filtered at 800 Hz using a Low-Pass Bessel Filter 8 Pole (Warner Instruments) and digitized at 4 kHz. All experiments were performed at room temperature (23 ± 2°C). The recordings were analyzed using Clampfit 10.6 (Molecular Devices) and Prism (ver. 5.0, GraphPad).

### Biochemical characterization of CHD of *Gm*QUAC1

cDNAs encoding the C-terminal cytoplasmic helical domain of *Gm*QUAC1 (wild-type or mutants) were subcloned into a modified bacterial expression vector pET-24a, with a C-terminal 10× His tag. Transformed *E. coli* BL21 (DE3) cells were grown in a TB medium containing Kanamycin (50 μg/ml). Protein expression was induced in cells grown to an optical density at OD600 of ~0.8 with 0.4 mM isopropyl β-D-thiogalactoside (IPTG) and followed by overnight cell growth at 16°C. Cells were collected and subsequently lysed using a high-pressure cell disrupter (JNBIO) with 3 passes at ~15,000 psi. The target protein was purified by using Ni^2+^ affinity chromatography and further analyzed by Superdex75 (10/300) gel filtration chromatography and SDS-PAGE.

### The three-dimensional structure modelling using AlphaFold2

Briefly, the sequence for AlphaFold prediction was generated by two individual *Gm*QUAC1 sequences, connecting by a linker of tandemly repeated sequence of (GSGS)_50_. Running with the local installed AlphaFold2 program and the CASP14 preset and databases (as of July 23, 2021), five dimer models were generated, with their pLDDT (predicted lDDT-Cα) scores ranging from 64.6 to 68.8. All the AlphaFold predicted structures (AF-structure) are nearly identical, with apparent two-fold symmetry. The two protomers in the top-rank dimer model (scored 68.8) display an r.m.s.d of 0.92 Å, was selected for further structural comparison with the cryo-EM structure. For a better view, the disordered (residues 1-35, 394-454 and 507-537) regions were removed in the final model.

## QUANTIFICATION AND STATISTICAL ANALYSIS

In all figure legends, n represents the number of independent biological replicates. All quantitative data were presented as mean ± SEM.

## DATA AND CODE AVAILABILITY

All data are available in the manuscript or the supplementary material. The accession number for the 3D cryo-EM density map reported in this paper is XXX, and the Protein Data Bank (PDB) accession code for the coordinate is XXX.

## ADDITIONAL RESOURCES

This study did not generate any additional resources.

