## Supplemental Material for "Molecular basis for the R-type anion channel QUAC1 activity in guard cells"

### Supplementary Information

(1) Extended Data Fig. 1-10
(2) Extended Data Table 1-2
(3) Supplementary Video 1-2

### Extended Data Fig. 1

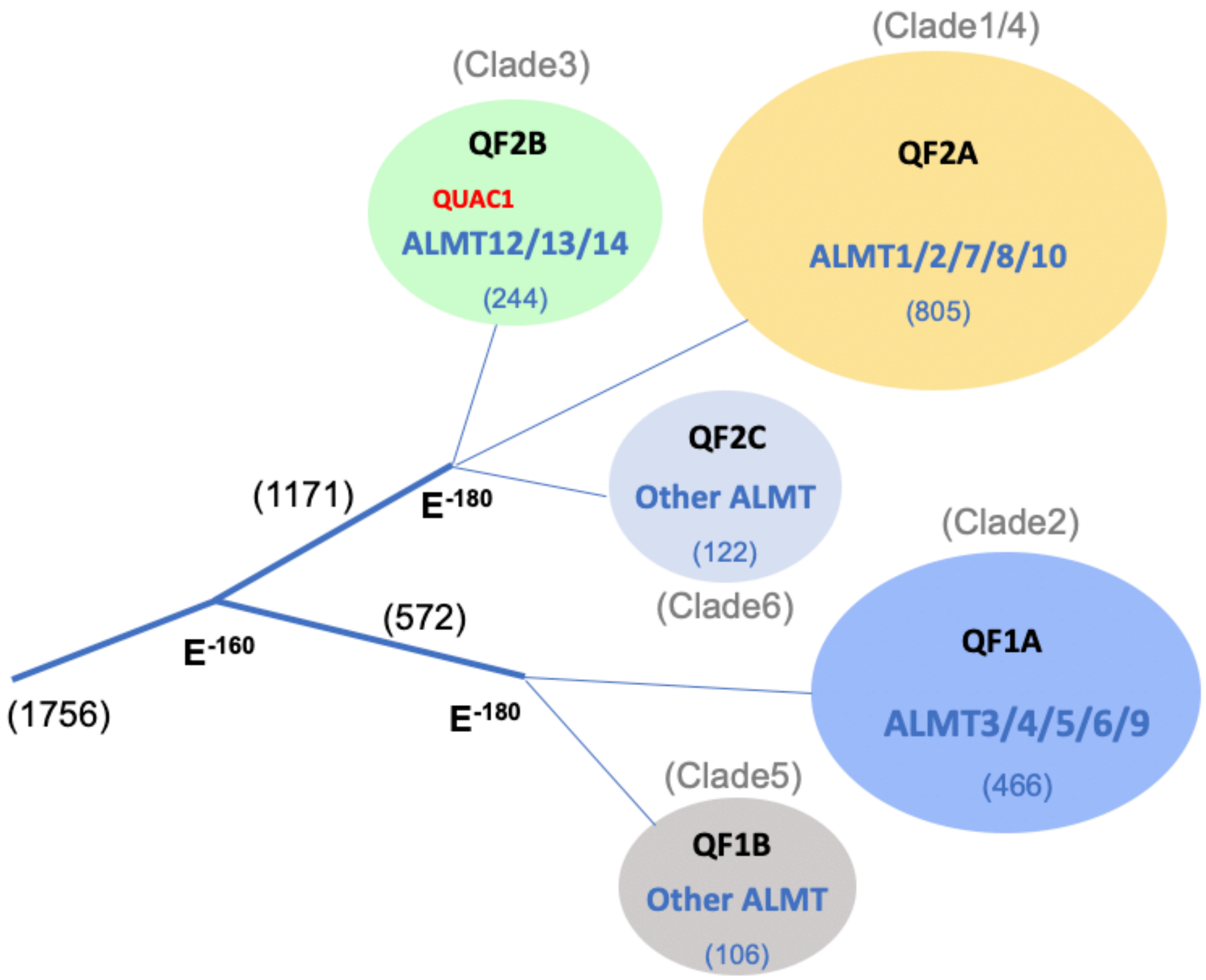

**Extended Data Fig. 1**

**Family tree of the QUAC/ALMTs**

The presentation was computed by the program COBALT[50] from representative sequences, including thirteen *Arabidopsis* ALMTs, and two other ALMTs. The clustering of sequences into a subfamily was performed using PSI-BLAST at the different levels ($E \leq 10^{-160}$, $E \leq 10^{-180}$) with representative sequences, as detailed in the Extended Data Table 1. The number in parentheses indicates the sequences in the family/subfamily. The clade numbering in previous work[51] is also indicated, and the Clade6 is a new group.

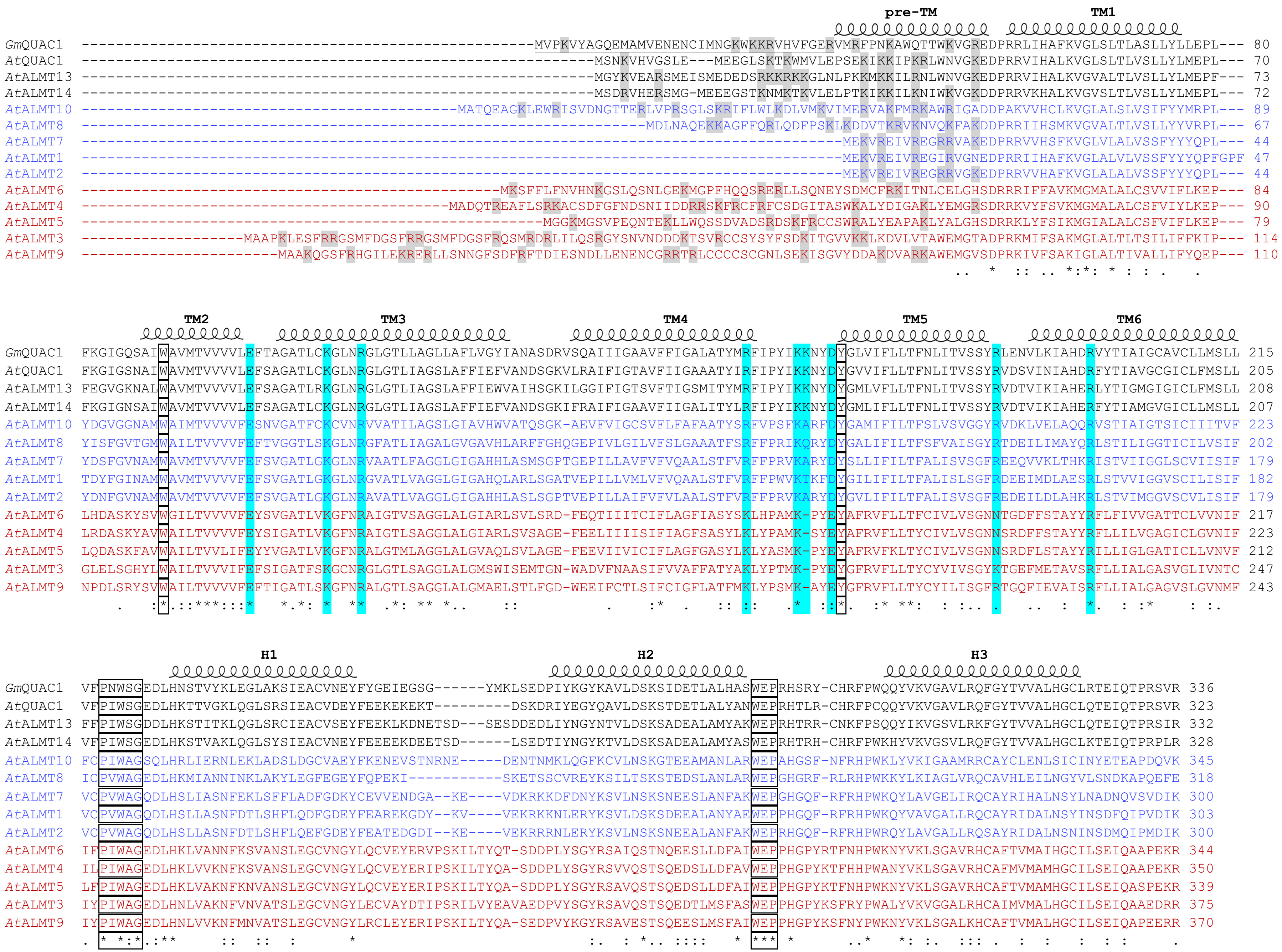

pre-TM
TM1
GmQUAC1 -----------------------------------------------------MVPKVYAGQEMAMVENENCIMNGKWKKRVHVFGERVMRFPNKAWQTTWKVGREDPRRLIHAFKVGLSLTLASLLYLLEPL--- 80
AtQUAC1 ------------------------------------------------------------MSNKVHVGSLE---MEEGLSKTKWMVLEPSEKIKKIPKRLWNVGKEDPRRVIHALKVGLSLTLVSLLYLMEPL--- 70
AtALMT13 ------------------------------------------------------------MGYKVEARSMEISMEDEDSRKKRKKGLNLPKKMKKILRNLWNVGKEDPRRVIHALKVGVALTLVSLLYLMEPF--- 73
AtALMT14 ------------------------------------------------------------MSDRVHERSMG-MEEEGSTKNMKTKVLELPTKIKKILKNIWKVGKDDPRRVKHALKVGVSLTLVSLLYLMEPL--- 72
AtALMT10 ----------------------------------------MATQEAGKLEWRISVDNGTTERLVPRSGLSKRIFLWLKDLVMKVIMERVAKFMRKAWRIGADDPAKVVHCLKVGLALSLVSIFYYMRPL--- 89
AtALMT8 ---------------------------------------------------------------MDLNAQEKKAGFFQRLQDFPSKLKDDVTKRVKNVQKFAKDDPRRIIHSMKVGVALTLVSLLYYVRPL--- 67
AtALMT7 ----------------------------------------------------------------------------------------MEKVREIVREGRRVAKEDPRRVVHSFKVGLVLALVSSFYYYQPL--- 44
AtALMT1 ----------------------------------------------------------------------------------------MEKVREIVREGIRVGNEDPRRIIHAFKVGLALVLVSSFYYYQPFGPF 47
AtALMT2 ----------------------------------------------------------------------------------------MEKVREIVREGRRVGKEDPRRVVHAFKVGLALALVSSFYYYQPL--- 44
AtALMT6 ------------------------------------------------MKSFFLFNVHNKGSLQSNLGEKMGPFHQQSRERLLSQNEYSDMCFRKITNLCELGHSDRRRIFFAVKMGMALALCSVVIFLKEP--- 84
AtALMT4 ------------------------------------------MADQTREAFLSRKACSDFGFNDSNIIDDRRSKFRCFRFCSDGITASWKALYDIGAKLYEMGRSDRRKVYFSVKMGMALALCSFVIYLKEP--- 90
AtALMT5 ----------------------------------------------------MGGKMGSVPEQNTEKLLWQSSDVADSRDSKFRCCSWRALYEAPAKLYALGHSDRRKLYFSIKMGIALALCSFVIFLKEP--- 79
AtALMT3 -------------------MAAPKLESFRRGSMFDGSFRRGSMFDGSFRQSMRDRLILQSRGYSNVNDDDKTSVRCCSYSYFSDKITGVVKKLKDVLVTAWEMGTADPRKMIFSAKMGLALTLTSILIFFKIP--- 114
AtALMT9 -----------------------MAAKQGSFRHGILEKRERLLSNNGFSDFRFTDIESNDLLENENCGRRTRLCCCCSCGNLSEKISGVYDDAKDVARKAWEMGVSDPRKIVFSAKIGLALTIVALLIFYQEP--- 110
.. * :: .. *:*: * : : . .
TM2
TM3
TM4
TM5
TM6
GmQUAC1 FKGIGQSAIWAVMTVVVVLEFTAGATLCKGLNRGLGTLLAGLLAFLVGYIANASDRVSQAIIIGAAVFFIGALATYMRFIPYIKKNYDYGLVIFLLTFNLITVSSYRLENVLKIAHDRVYTIAIGCAVCLLMSLL 215
AtQUAC1 FKGIGSNAIWAVMTVVVVLEFSAGATLCKGLNRGLGTLIAGSLAFFIEFVANDSGKVLRAIFIGTAVFIIGAAATYIRFIPYIKKNYDYGVVIFLLTFNLITVSSYRVDSVINIAHDRFYTIAVGCGICLFMSLL 205
AtALMT13 FEGVGKNALWAVMTVVVVLEFSAGATLRKGLNRGLGTLIAGSLAFFIEWVAIHSGKILGGIFIGTSVFTIGSMITYMRFIPYIKKNYDYGMLVFLLTFNLITVSSYRVDTVIKIAHERLYTIGMGIGICLFMSLL 208
AtALMT14 FKGIGNSAIWAVMTVVVVLEFSAGATLCKGLNRGLGTLIAGSLAFFIEFVANDSGKIFRAIFIGAAVFIIGALITYLRFIPYIKKNYDYGMLIFLLTFNLITVSSYRVDTVIKIAHERFYTIAMGVGICLLMSLL 207
AtALMT10 YDGVGGNAMWAIMTVVVVFESNVGATFCKCVNRVVATILAGSLGIAVHWVATQSGK-AEVFVIGCSVFLFAFAATYSRFVPSFKARFDYGAMIFILTFSLVSVGGYRVDKLVELAQQRVSTIAIGTSICIIITVF 223
AtALMT8 YISFGVTGMWAILTVVVVFEFTVGGTLSKGLNRGFATLIAGALGVGAVHLARFFGHQGEPIVLGILVFSLGAAATFSRFFPRIKQRYDYGALIFILTFSFVAISGYRTDEILIMAYQRLSTILIGGTICILVSIF 202
AtALMT7 YDSFGVNAMWAVMTVVVVFEFSVGATLGKGLNRVAATLFAGGLGIGAHHLASMSGPTGEPILLAVFVFVQAALSTFVRFFPRVKARYDYSLLIFILTFALISVSGFREEQVVKLTHKRISTVIIGGLSCVIISIF 179
AtALMT1 TDYFGINAMWAVMTVVVVFEFSVGATLGKGLNRGVATLVAGGLGIGAHQLARLSGATVEPILLVMLVFVQAALSTFVRFFPWVKTKFDYGILIFILTFALISLSGFRDEEIMDLAESRLSTVVIGGVSCILISIF 182
AtALMT2 YDNFGVNAMWAVMTVVVVFEFSVGATLGKGLNRAVATLVAGGLGIGAHHLASLSGPTVEPILLAIFVFVLAALSTFVRFFPRVKARYDYGVLIFILTFALISVSGFREDEILDLAHKRLSTVIMGGVSCVLISIF 179
AtALMT6 LHDASKYSVWGILTVVVVFEYSVGATLVKGFNRAIGTVSAGGLALGIARLSVLSRD-FEQTIIITCIFLAGFIASYSKLHPAMK-PYEYAFRVFLLTFCIVLVSGNNTGDFFSTAYYRFLFIVVGATTCLVVNIF 217
AtALMT4 LRDASKYAVWAILTVVVVFEYSIGATLVKGFNRAIGTLSAGGLALGIARLSVSAGE-FEELIIIISIFIAGFSASYLKLYPAMK-SYEYAFRVFLLTYCIVLVSGNNSRDFFSTAYYRFLLILVGAGICLGVNIF 223
AtALMT5 LQDASKFAVWAILTVVLIFEYYVGATLVKGFNRALGTMLAGGLALGVAQLSVLAGE-FEEVIIVICIFLAGFGASYLKLYASMK-PYEYAFRVFKLTYCIVLVSGNNSRDFLSTAYYRILLIGLGATICLLVNVF 212
AtALMT3 GLELSGHYLWAILTVVVIFEFSIGATFSKGCNRGLGTLSAGGLALGMSWISEMTGN-WADVFNAASIFVVAFFATYAKLYPTMK-PYEYGFRVFLLTYCYVIVSGYKTGEFMETAVSRFLLIALGASVGLIVNTC 247
AtALMT9 NPDLSRYSVWAILTVVVVFEFTIGATLSKGFNRALGTLSAGGLALGMAELSTLFGD-WEEIFCTLSIFCIGFLATFMKLYPSMK-AYEYGFRVFLLTYCYILISGFRTGQFIEVAISRFLLIALGAGVSLGVNMF 243
. :*.::***:::* *.*: * ** .*: ** *.. :: . :* . :: :: .* ::*. :* **: : :.. . .. : *. : :* : :.
H1
H2
H3
GmQUAC1 VFPNWSGEDLHNSTVYKLEGLAKSIEACVNEYFYGEIEGSG------YMKLSEDPIYKGYKAVLDSKSIDETLALHASWEPRHSRY-CHRFPWQQYVKVGAVLRQFGYTVVALHGCLRTEIQTPRSVR 336
AtQUAC1 VFPIWSGEDLHKTTVGKLQGLSRSIEACVDEYFEEKEKEKT---------DSKDRIYEGYQAVLDSKSTDETLALYANWEPRHTLR-CHRFPCQQYVKVGAVLRQFGYTVVALHGCLQTEIQTPRSVR 323
AtALMT13 FFPIWSGDDLHKSTITKLQGLSRCIEACVSEYFEEKLKDNETSD---SESDDEDLIYNGYNTVLDSKSADEALAMYAKWEPRHTRR-CNKFPSQQYIKVGSVLRKFGYTVVALHGCLQTEIQTPRSIR 332
AtALMT14 VFPIWSGEDLHKSTVAKLQGLSYSIEACVNEYFEEEEKDEETSD------LSEDTIYNGYKTVLDSKSADEALAMYASWEPRHTRH-CHRFPWKHYVKVGSVLRQFGYTVVALHGCLKTEIQTPRPLR 328
AtALMT10 FCPIWAGSQLHRLIERNLEKLADSLDGCVAEYFKENEVSTNRNE-----DENTNMKLQGFKCVLNSKGTEEAMANLARWEPAHGSF-NFRHPWKLYVKIGAAMRRCAYCLENLSICINYETEAPDQVK 345
AtALMT8 ICPVWAGEDLHKMIANNINKLAKYLEGFEGEYFQPEKI-----------SKETSSCVREYKSILTSKSTEDSLANLARWEPGHGRF-RLRHPWKKYLKIAGLVRQCAVHLEILNGYVLSNDKAPQEFE 318
AtALMT7 VCPVWAGQDLHSLIASNFEKLSFFLADFGDKYCEVVENDGA--KE----VDKRKKDFDNYKSVLNSKSNEESLANFAKWEPGHGQF-RFRHPWKQYLAVGELIRQCAYRIHALNSYLNADNQVSVDIK 300
AtALMT1 VCPVWAGQDLHSLLASNFDTLSHFLQDFGDEYFEAREKGDY--KV----VEKRKKNLERYKSVLDSKSDEEALANYAEWEPPHGQF-RFRHPWKQYVAVGALLRQCAYRIDALNSYINSDFQIPVDIK 303
AtALMT2 VCPVWAGQDLHSLLASNFDTLSHFLQEFGDEYFEATEDGDI--KE----VEKRRRNLERYKSVLNSKSNEEALANFAKWEPRHGQF-RFRHPWRQYLAVGALLRQSAYRIDALNSNINSDMQIPMDIK 300
AtALMT6 IFPIWAGEDLHKLVANNFKSVANSLEGCVNGYLQCVEYERVPSKILTYQT-SDDPLYSGYRSAIQSTNQEESLLDFAIWEPPHGPYRTFNHPWKNYVKLSGAVRHCAFTVMAIHGCILSEIQAAPEKR 344
AtALMT4 ILPIWAGEDLHKLVVKNFKSVANSLEGCVNGYLQCVEYERIPSKILTYQA-SDDPLYSGYRSVVQSTSQEDSLLDFAVWEPPHGPYKTFHHPWANYVKLSGAVRHCAFMVMAMHGCILSEIQAAPEKR 350
AtALMT5 LFPIWAGEDLHKLVAKNFKNVANSLEGCVNGYLQCVEYERIPSKILTYQA-SDDPLYSGYRSAVQSTSQEDSLLDFAIWEPPHGPYKTFNHPWKNYVKLSGAVRHCAFTVMAMHGCILSEIQASPEKR 339
AtALMT3 IYPIWAGEDLHNLVAKNFVNVATSLEGCVNGYLECVAYDTIPSRILVYEAVAEDPVYSGYRSAVQSTSQEDTLMSFASWEPPHGPYKSFRYPWALYVKVGGALRHCAIMVMALHGCILSEIQAAEDRR 375
AtALMT9 IYPIWAGEDLHNLVVKNFMNVATSLEGCVNGYLRCLEYERIPSKILTYQA-SEDPVYKGYRSAVESTSQEESLMSFAIWEPPHGPYKSFNYPWKNYVKLSGALKHCAFTVMALHGCILSEIQAPEERR 370
. * *:*.:** :: :: : * :. : *.. :::: * *** * ..* *: :. ::: . : : : : : .

```
                        H4                                 H5
GmQUAC1   AMFKDPCIRLAAEVSKVLIELSNSIRNRRHCS--PEILSDHLHEALQDLNTAIKSQPRLFLGPKHRHNQATNMLKIA-----------------------------------------AAQVGQERHGKTSLS 426
AtQUAC1   ALFKDPCVRLAGEVCKALTELADSISNHRHCS--PEILSDHLHVALQDLNSAIKSQPKLFLGSNLHRHNNKHQNGSISNNKHHQRNSSNSGKDLNGDVSLQNTETGTRKIT-------ETGSRQGQNGAVSLS 447
AtALMT13  VLFKDPCVRLAGEICKVLSELSESIQNRRHCS--SEILSDSLEAALKDLNSTIKSQPKLFLGSNLHSNITNKHLNGH--------VSYYNETNSNGTVSYHNDNNTNGCVLGETIEENDTVSPLPLNSVVSLS 455
AtALMT14  GLFKDPCVRLAGEICKVLSELAASIRNRRHCS--PEILSDSLQVALQDLNTAIKSQPKLFLGSSQNGNVSQGNSGRHNPNV---AVSQHINKDTNEAASYQNTGTPRG----------ERMSRFGP--NVSFS 444
AtALMT10  NHFGEACMKLSSASSKILRELADMMKNTRKSS-KMDFLVFDMNSAVQELQETLKNVPIETNKPEEVPSEEENK------------------------------------------------------------ 413
AtALMT8   SKIQEPITTMSREVGEALKAIAKSIKTMRNDSACVNAHIDNSKKAIKNLKIALKSSYPETY------------------------------------------------------------------------ 379
AtALMT7   KKLGEPLRRMSLESGKAMKEMSISLKKMTKPS-SSDLHVQNAKSACKSLTNLLNSGILKE------------------------------------------------------------------------- 359
AtALMT1   KKLETPLRRMSSESGNSMKEMSISLKQMIKSS-SSDIHVSNSQAACKSLSTLLKSGILND------------------------------------------------------------------------- 362
AtALMT2   KKIEEPLRRMSSESGKSMKEVSISLKNMTISS-SFDIHVVNSQSACKTLSTLLKSGILND------------------------------------------------------------------------- 359
AtALMT6   QAFRHELQRVGNEGAKVLRLIGEKVEKMENL--GPGEILNDVQRAAEELQMKIDSKSYLLVNSESWAATKEKAEAEEYEEE-------------------AHETKVIKSLSQIWDTNSSSNNQNPASG----- 451
AtALMT4   QAFRQELQRVGNEGAKVLRLFGEKVEKMEKLS--PGNVLKDVQRAAEELQMKIDSNSFLLVNSESWAAMKEKAEAEEAQQN--------------YHEAKDDESKVIQSLSQIWDNNNNPHHQNQHA------ 461
AtALMT5   HVFSNELRRVGNEGAKVLRLFGEKVEKMEKLSLSLGEILKDVQRAAEALQMKIDSKSYLLVNSESWAAIKEQAEAEEAREN--------------DQEAKDDETKVIKSLSQIWDTNNNNNHQS--------- 449
AtALMT3   REFRNELQRVGIEGAKVLRYIGESLKKMEKLN-PIEDILYEIHQAAEELQSKIDKKSYLLVNAKNWEIGNRPRVRDLTDEQ----KISNLDSDLSRILAHKSQSEATLRPPKNWDDVTTAANLSSA------- 496
AtALMT9   QVFRQELQRVGVEGAKLLRELGEKVKKMEKLG-PV-DLLFEVHLAAEELQHKIDKKSYLLVNSECWEIGNRATKESEPQELLSLEDSDPPENHAPPIYAFKSLSEAVLEIPPSWGEKNHREA----------- 490
            :      :.    : :  ..  :                 . * : *   :..

                                                                     H6                      H7
GmQUAC1   SVKTDSSALLEWKTKR--VSAEQTKESERKSLRP-------------------QLSKIA-ITSLEFSEALPFAAFASLLVETVAKLDLVIEEVEELGRLACFKEFIPGDEFVVTCQEPRVDVSQNHLPSHGVD 537
AtQUAC1   SFRTDTSALMEYRRSFKNSNSEMSAAGERRMLRP-------------------QLSKIAVMTSLEFSEALPFAAFASLLVEMVARLDNVIEEVEELGRIASFKEYDNKRDQTADDVRCENPANVTISVGAAE- 560
AtALMT13  SLRS---------------VKKSAATGEKRRLRK-------------------QLSKIAVMKSLEFSEALPFAAFASLLVEMVARLDTVIDEVEELGTIACFKEYDKTVEVRIENRLI--------------- 539
AtALMT14  RLRADT-----------LERRSAAATNERKILRQ-------------------QLSRIVVLTSLEFSEALPFAAFASLLVEMVARLDNVIEEVEELGTIACFKDYDNNVDQKDVEVRVEKPADLVVGVE---- 543
AtALMT10  ------------------------------------------------------VDSEERTTSMSLHEVLPVATLVSLLIENAARIQTAVEAVDELANLADFEQDSKKKTGDNNTKQPPLSS----------- 485
AtALMT8   ---------------------------------------------------------------KDLLEIIPGVTMASILIEVVNCVEKIYEAVEEFSGLAHFKETLDSKLSAEIGQHQLLHRGCVKPVLDGDN 446
AtALMT7   ---------------------------------------------------------------VEPLELVSLLTAISLLIDIINLTEKILESLHELATAAKFKNKIEHPLFSEKPKAKSFVSVRSIKCHDDHV 426
AtALMT1   ---------------------------------------------------------------VEPLQMISLMTTVSMLIDIVNLTEKISESVHELASAARFKNKMRPTVLYEKSDSGSIGRAMPIDSHEDHH 429
AtALMT2   ---------------------------------------------------------------VEPLQMISLMTTVSLLIDIVNLTEKISESVHELASAAKFKNKKKPS----KSNSGSIGQAMPNKSHDDDD 422
AtALMT6   -----------------NDESQIWESTESMMLRNRETWPSVSFI-------GGSVVNETVYKVYESASSLSLATFASLLIEFVARLENLVNAFEELSTKADFRDPVPLNVVDQEGLWTTLMRLRRLRRYFGRS 560
AtALMT4   -----------------GNDSQLWISTESMMLRNRENWPSVSFI-------GGSMINEIESKVYESASSLSLATFASLLIEFVARLQNIVNAYEELSTKADFKEQVSETRI---------------------- 548
AtALMT5   -----------------NDQSQHWMSTESMMLKNREMWPSMSFI-------DGTVVNEIECKVYESASSLSLATFASLLIEFVARLQNIVNAFEELSTKAGFKDAVDQIPKV--------------------- 537
AtALMT3   --------------------TMLPYLQSRTMIHKQPSWPSRISI---TPGSMLQPPLGEPGKMYESASNLSLATFASLLIEFVARLENLVNAYDELSVKANFKEAVSE------------------------- 581
AtALMT9   -------------------------LNHRPTFSKQVSWPARLVLPPHLETTNGASPLVETTKTYESASALSLATFASLLIEFVARLQNVVDAFKELSQKANFKEPEIVTTGTDVEFSGERVGLGQKIRRCFGM 598
                                                                          .  . :   :  *:*::     :   :  .*:.  * *.:

GmQUAC1   ---------------------------------------------------------------------------- 537
AtQUAC1   ---------------------------------------------------------------------------- 560
AtALMT13  ---------------------------------------------------------------------------- 539
AtALMT14  ---------------------------------------------------------------------------- 543
AtALMT10  ---------------------------------------------------------------------------- 485
AtALMT8   EKEDNSSCHVLITVHDEGYLPTATAKNVLGAEKTRVDIV------------------------------------- 488
AtALMT7   VIIIEDDGNNDDTSKNDNGSKEVSIHEKHEDDDTHVDARCVSCGHTSVCVK------------------------- 480
AtALMT1   VVTVLHDVDNDRSNNVDDSRGGSSQDSCHHVAIKIVDDNSNHEKHEDGEIHVHTLSNGHLQ--------------- 493
AtALMT2   HVVTILGDVDTSNNVDQSQSHGEISVDSCHHVTIKINDDDSIHDKNEDGDIHVHTNRVSCDHTNASDLLDSGVKKN 501
AtALMT6   ---------------------------------------------------------------------------- 560
AtALMT4   ---------------------------------------------------------------------------- 548
AtALMT5   ---------------------------------------------------------------------------- 537
AtALMT3   ---------------------------------------------------------------------------- 581
AtALMT9   ---------------------------------------------------------------------------- 598
```

**Extended Data Fig. 2**

**Structure-based sequence alignment for plant QUAC/ALMTs**

Structure-based sequence alignments of thirteen *Arabidopsis* ALMT/QUACs and *Glycine max* QUAC1. The structure of *Gm*QUAC1 has been used to restrict sequence gaps to inter-helical segments, with the superior coils defining extents of the helical segments. The disordered regions in *Gm*QUAC1 (residues 1-35, 394-454 and 507-537) are indicated by underlines. The aligned sequences are as following: *Gm*QUAC1 (XP_003516635.1), *At*QUAC1/ALMT12 (NP_193531.1), *At*ALMT1 (NP_172319.1), *At*ALMT2 (NP_172320.1), *At*ALMT3 (NP_173278.1), *At*ALMT4 (NP_173919.1), *At*ALMT5( NP_564935.1), *At*ALMT6 (NP_179338.1), *At*ALMT7 (NP_001324626.1), *At*ALMT8 (NP_187774.1), *At*ALMT9 (NP_188473.1), *At*ALMT10 (NP_001319836.1), *At*ALMT13 (NP_199472.1), *At*ALMT14 (NP_199473.1).

The protein sequences are from three major groups: QF1A (*At*ALMT3/4/5/6/9) in red; QF2A (*At*ALMT1/2/7/8/10) in blue; QF2B (*Gm*QUAC1/ *At*QUAC1/*At*ALMT13/14) in black. Overall, the protein sequences in the QF2A have a shorter pre-TM region, and lack a disordered region and a domain-swapped helix found in the *Gm*QUAC1 structure.

Some key residues are highlighted or indicated as following:

1. The pore-lining positively charged R/K residues are highlighted in cyan (K109/R113/R158/K164/K165/R187/R198 in *Gm*QUAC1), and related interacting negatively-charged E/D residues are also highlighted in cyan (E100/D168 in *Gm*QUAC1).
2. The residues of mutagenesis for disrupting dimer interaction are highlighted in yellow (S461/F470/L474/A477 in *Gm*QUAC1).
3. Positively charged residues at the N-terminal pre-TM juxtamembrane helix are highlighted in grey.
4. Potential phosphorylation sites in the disordered regions of C-terminal CHD are highlighted in green.
5. The domain-swapped helix regions are indicated in a box (H6 in *Gm*QUAC1).
6. Other conserved motifs are indicated in boxes (W90, Y169, P218-N219-W220-S221-G222 and W288-E289-P290 in *Gm*QUAC1).

### Extended Data Fig. 3

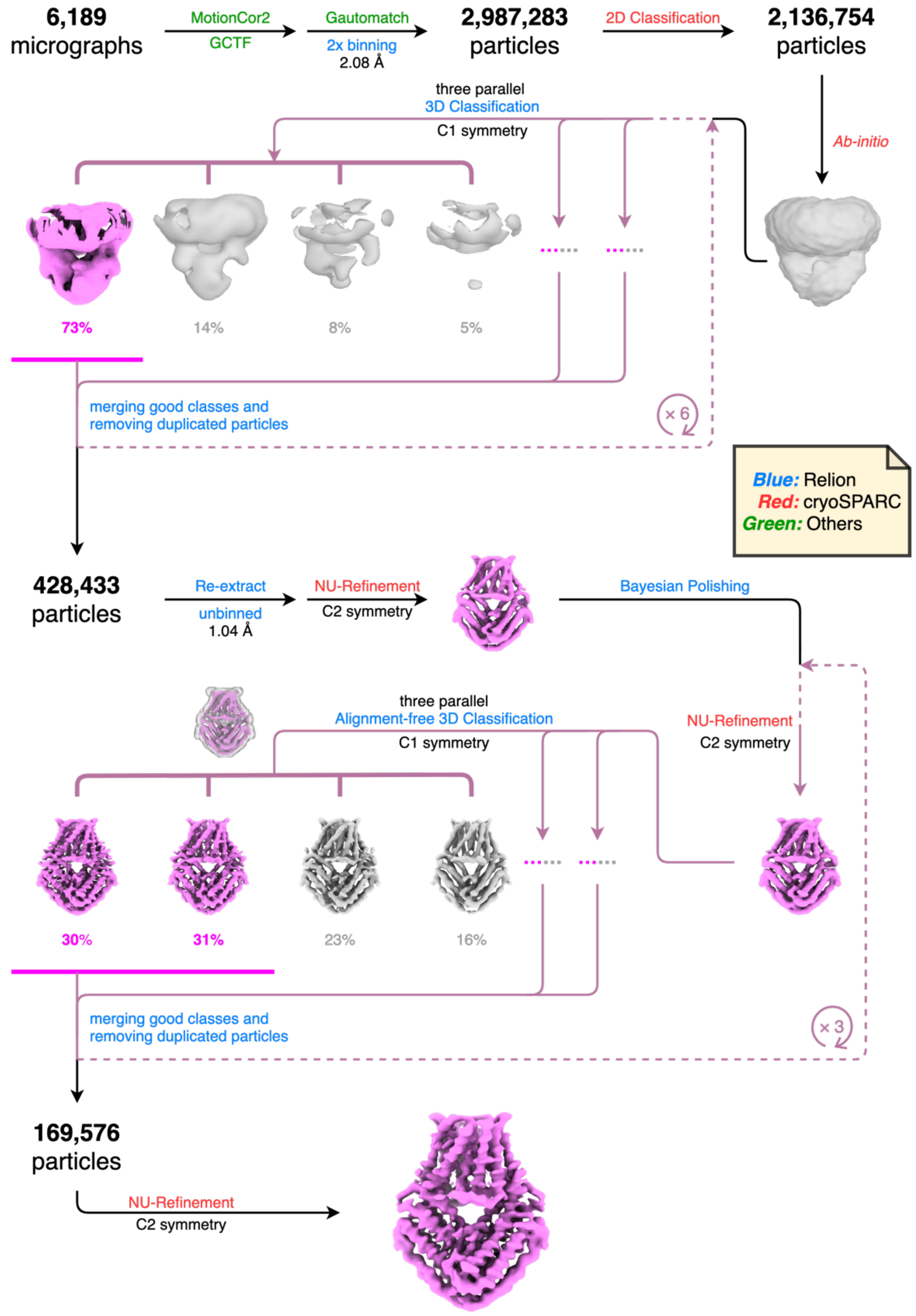

**Extended Data Fig. 3**

**The workflow for image processing of QUAC1**

### Extended Data Fig. 4

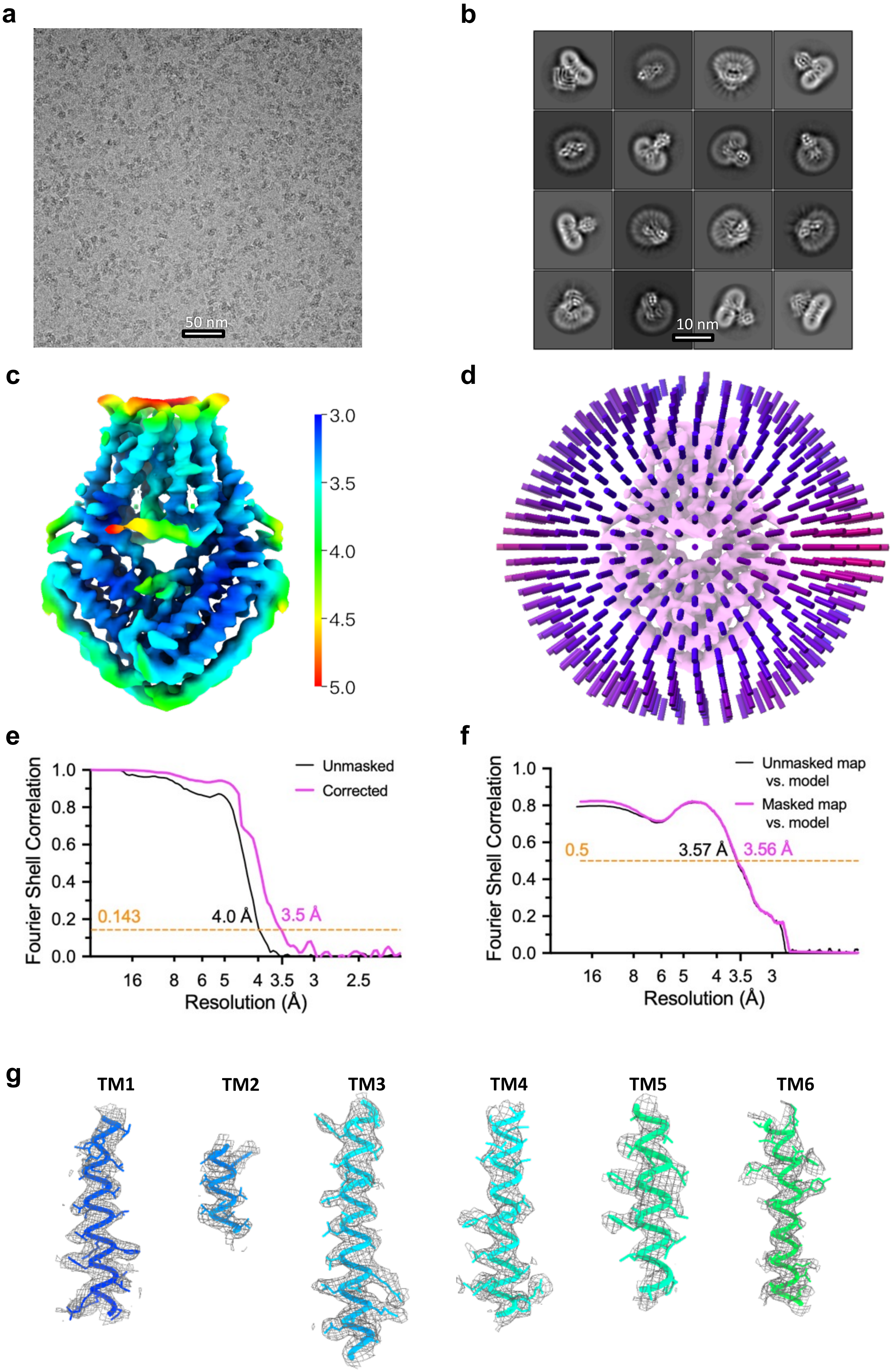

**Extended Data Fig. 4**

**Cryo-EM analysis of QUAC1**

**a,** Representative micrograph of *Gm*QUAC1.

**b,** Representative 2D class averages of *Gm*QUAC1.

**c,** Local resolution electron density map of *Gm*QUAC1

**d,** Euler angle distribution plot of particle projections in reconstruction of *Gm*QUAC1.

**e,** Fourier shell correlation (FSC) curve suggests an overall resolution at 3.5 Å, as estimated using the 0.143 cut-off criterion (dotted line).

**f,** Cross-validation of model to cryo-EM density map suggests a resolution at 3.5 Å, as estimated using the 0.5 cut-off criterion (dotted line).

All line charts in (**e**) and (**f**) were prepared in GraphPad Prism.

**g,** Representative cryo-EM density map for the TM segments.

### Extended Data Fig. 5

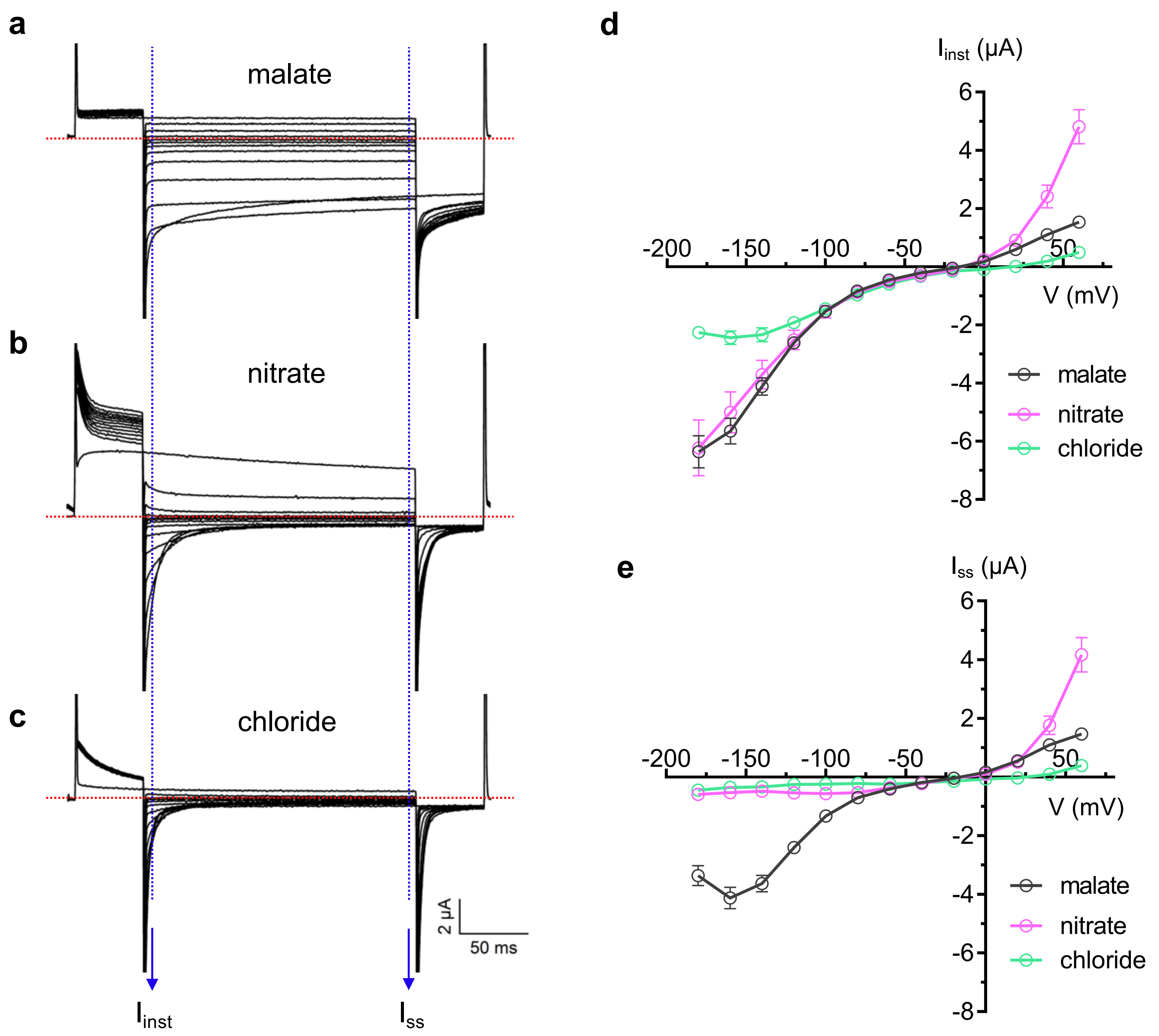

**Extended Data Fig. 5**

**TEVC recording of the QUAC1 channel in *Xenopus laevis* oocytes**

**a-c,** The representative current traces recorded at difference voltages (from +60 mV to –180 mV in 20 mV decrement) in various external solutions: 30 mM L-malate (**a**), 30 mM $NaNO_3$ (**b**) and 30 mM NaCl (**c**).

**d,** Instant current-voltage ($I_{inst}$-V) relations of *Gm*QUAC1 (Data are mean ± SEM, n≥8).

**e,** Steady-state current-voltage ($I_{ss}$-V) relations of *Gm*QUAC1 (Data are mean ± SEM, n≥8).

### Extended Data Fig. 6

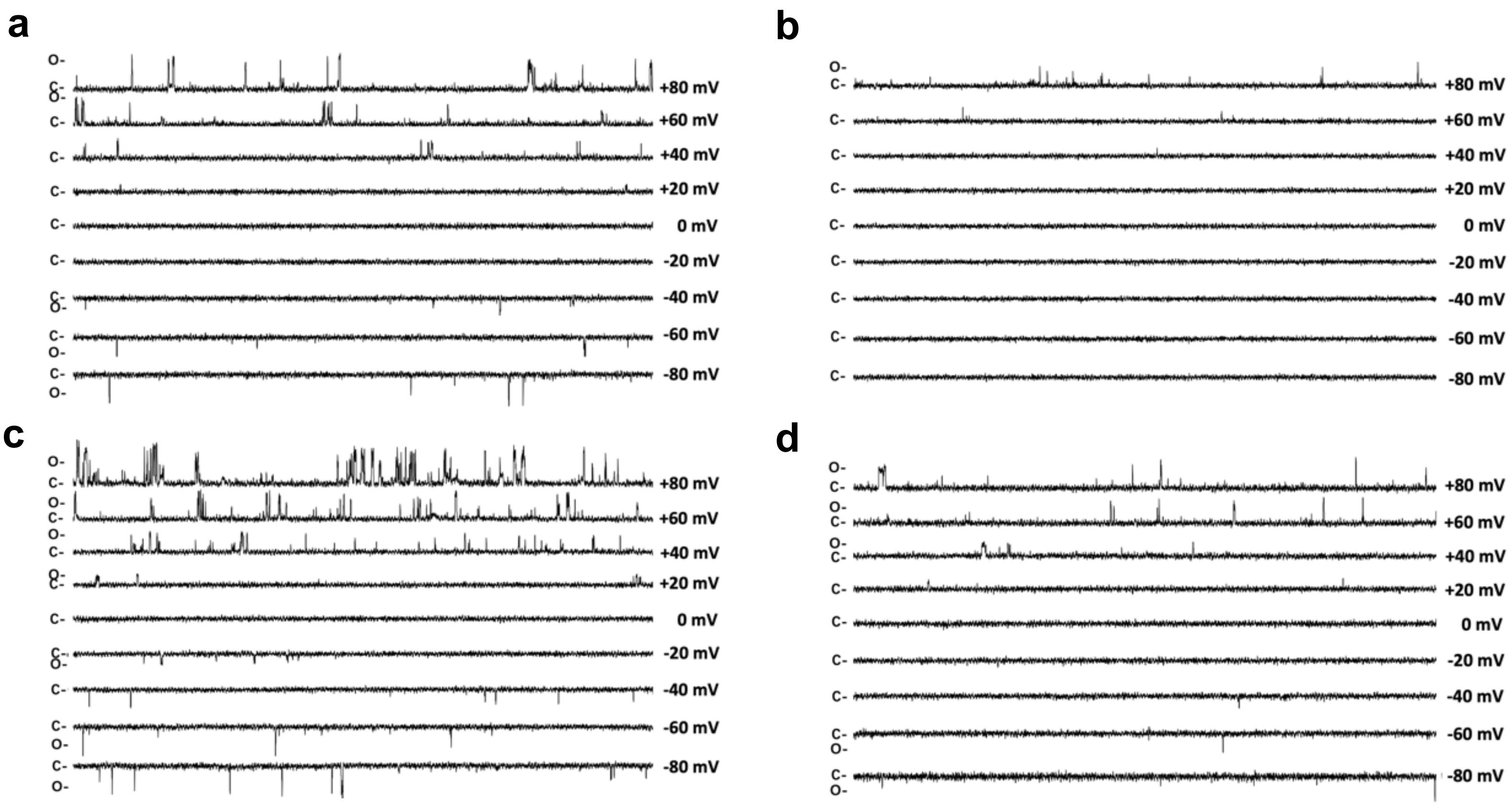

**Extended Data Fig. 6**

**Representative current traces for single channel analysis of QUAC1 channel in planar lipid bilayer**

**a**-**d**, Representative current traces for single channel analysis of *Gm*QUAC1 are shown at different holding potentials, as indicated. *Gm*QUAC1$^{NaCl}$ was tested in $NaNO_3$ (**a**) or NaCl (**b**) solution. *Gm*QUAC1$^{malate}$ was tested in $NaNO_3$ (**c**) or NaCl (**d**) solution.

### Extended Data Fig. 7

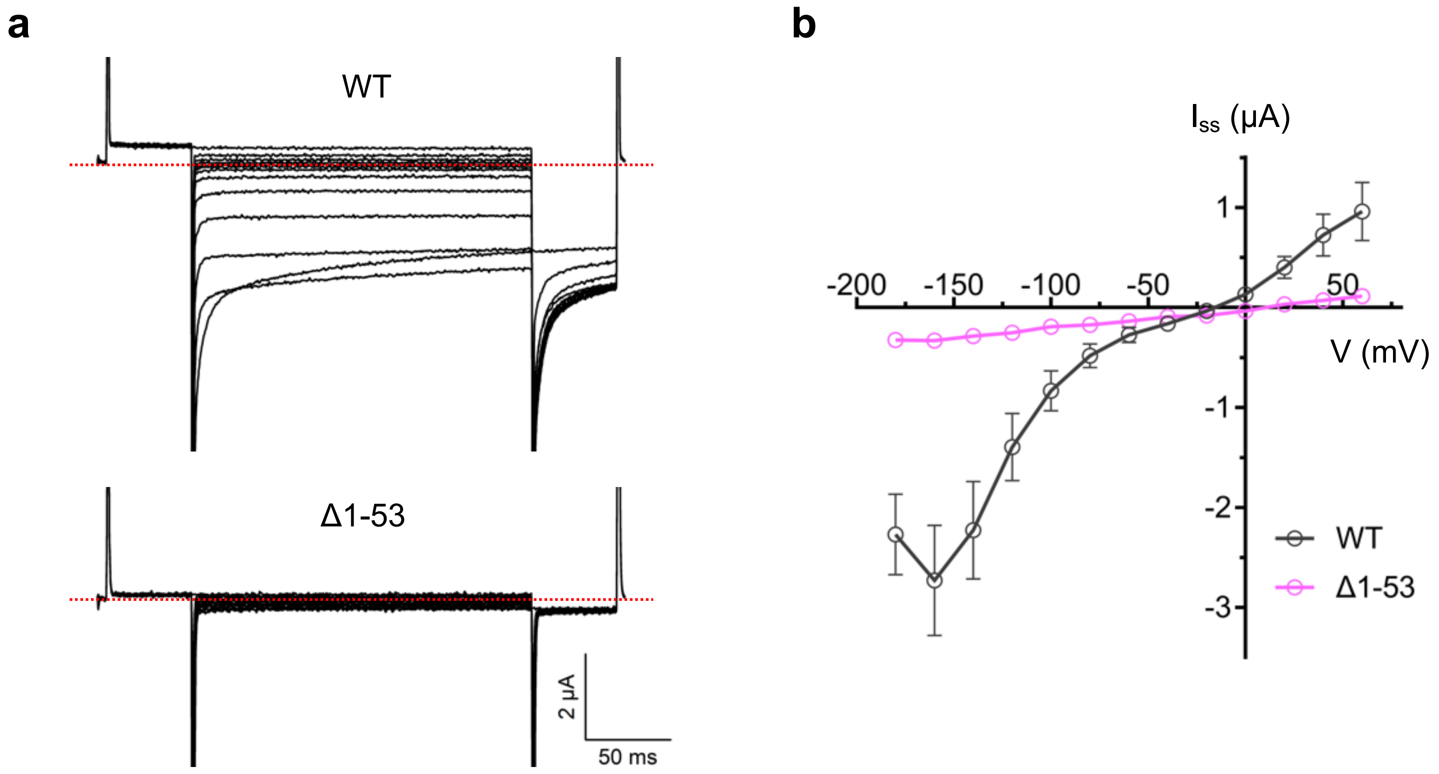

**Extended Data Fig. 7**

**Analysis of the QUAC1 pre-TM helix deletion mutant by TEVC recording, related to Figure S9**

**a,** Representative current traces recorded at different voltages (from +60 mV to –180 mV in 20 mV decrement) in the external solutions of 30 mM L-malate for the *Gm*QUAC1 wild-type (upper) and Δ1-53 mutant (lower).

**b,** Steady-state current-voltage ($I_{ss}$-V) relations of the *Gm*QUAC1 wild-type and Δ1-53 mutant (Data are mean ± SEM, n≥6).

### Extended Data Fig. 8

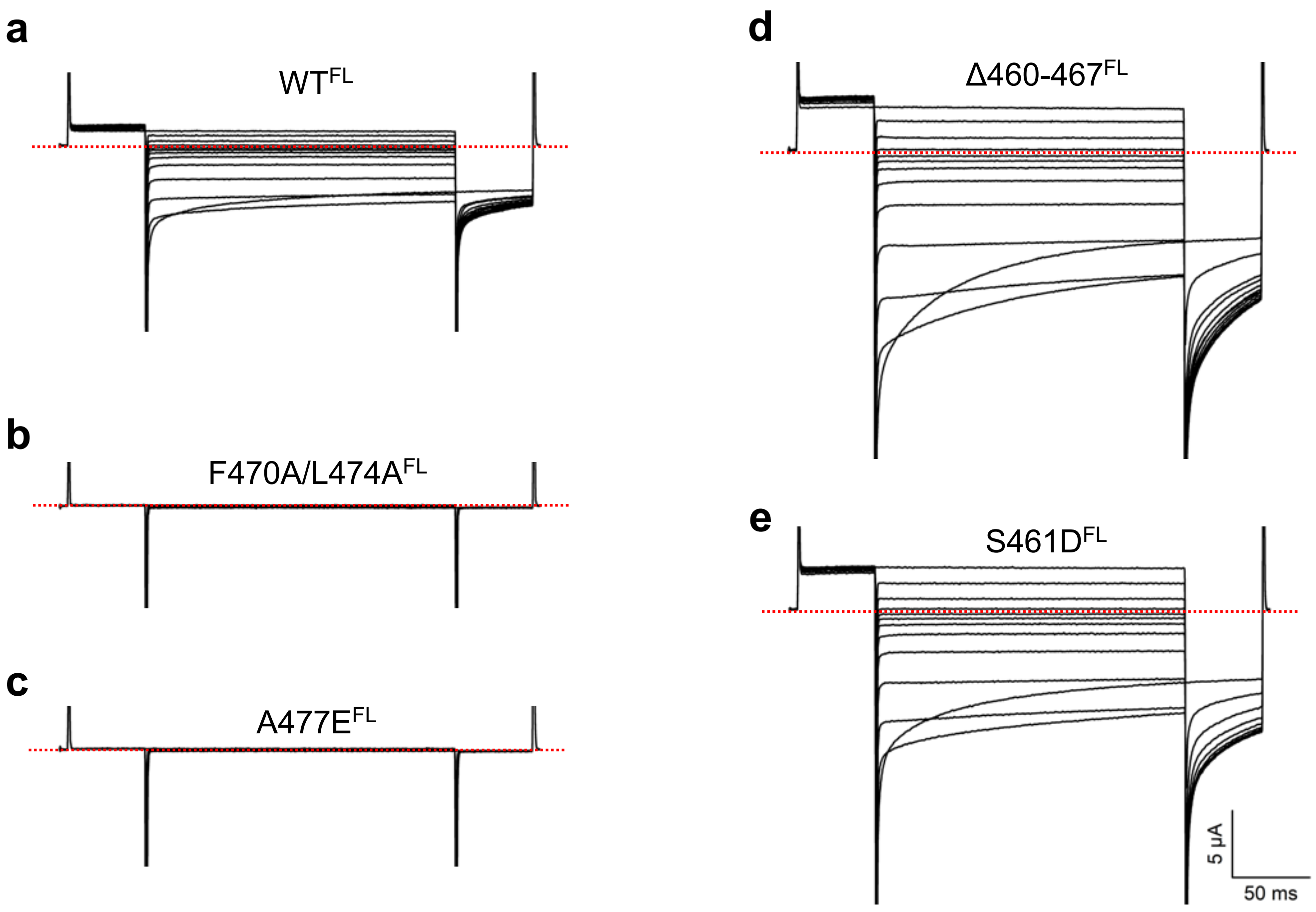

**Extended Data Fig. 8**

**Representative current traces of QUAC1 dimer-disrupting mutants by TEVC recording**

**a-e,** Representative current traces recorded at difference voltages (from +60 mV to –180 mV in 20 mV decrement) in the external solutions of 30 mM L-malate for the full-length *Gm*QUAC1 wild-type (**a**) and mutants: F470A/L474A (**b**), A477E (**c**), Δ460-467 (**d**) and S461D (**e**).

### Extended Data Fig. 9

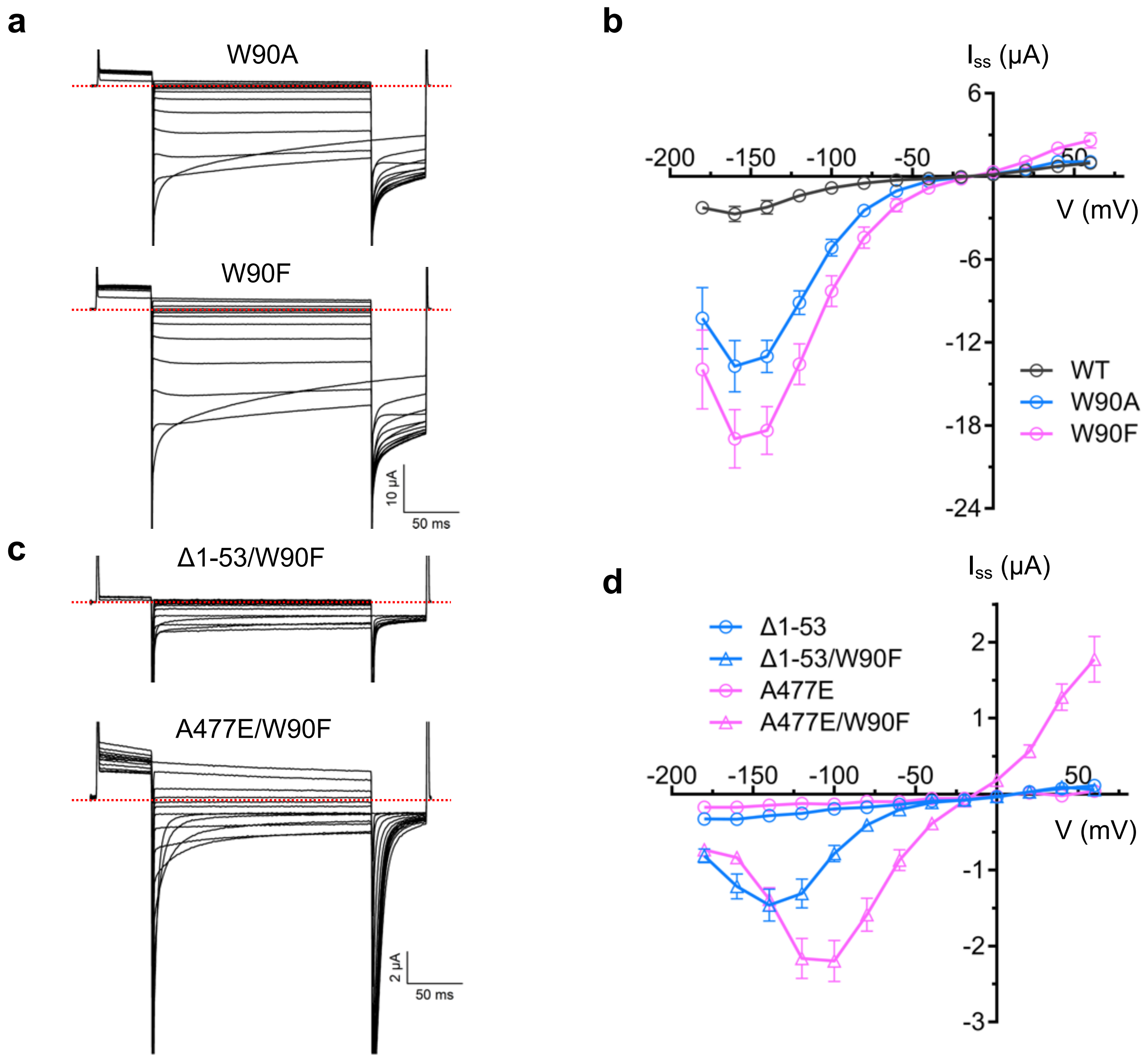

**Extended Data Fig. 9**

**Analysis of QUAC1 W90 mutants by TEVC recording**

**a** and **b,** The representative current traces (**a**) and steady-state current-voltage ($I_{ss}$-V) relations (**b**) of *Gm*QUAC1 mutants, W90A and W90F, at different applied voltages (from +60 mV to –180 mV in 20 mV decrement) in the external solutions of 30 mM L-malate (Data are mean ± SEM, n≥6).

**c** and **d,** The representative current traces (**c**) and steady-state current-voltage ($I_{ss}$-V) relations (**d**) of *Gm*QUAC1 mutants, Δ1-53/W90F and A477E/W90F, at different applied voltages (from +60 mV to –180 mV in 20 mV decrement) in the external solutions of 30 mM L-malate (Data are mean ± SEM, n≥6).

### Extended Data Fig. 10

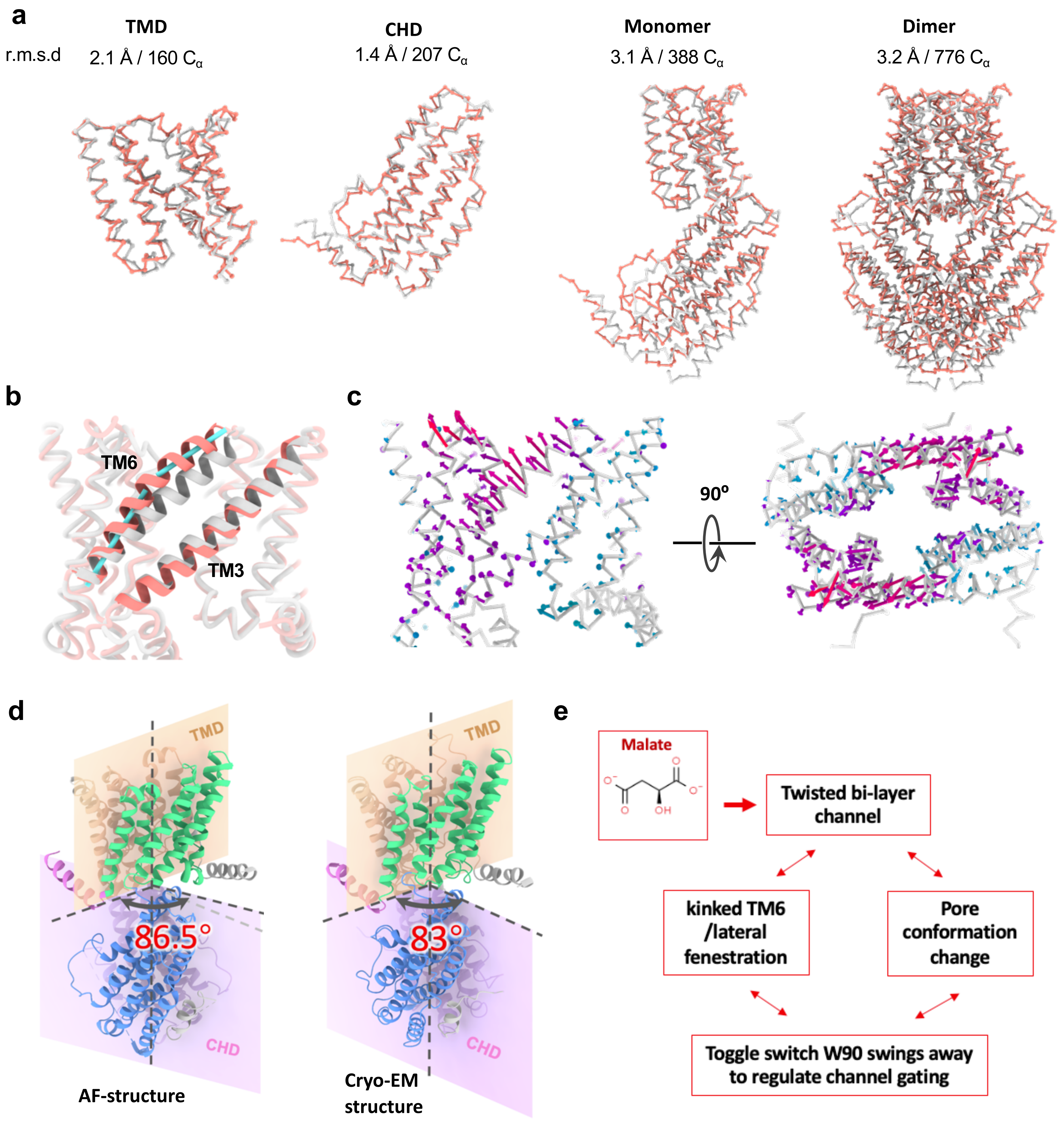

**Extended Data Fig. 10**

**Comparison of the cryo-EM structure and AF-structure**

**a,** Superimposition of the cryo-EM structure (salmon) and AF-structure (grey). The superimposed Cα and their r.m.s.d values are indicated.

**b,** The comparison of TMs in the cryo-EM structure (salmon) and AF-structure (grey). The TM3 and TM6 helices are shown in cartoon.

**c,** The porcupine plot showing the motion across the AF-structure and the cryo-EM structure. The arrows indicate direction of motion and magnitude of the movements of the Cα positions (from the AF-structure to the cryo-EM structure). The Cα traces for the AF-structure is shown. The colors for the arrows are indicated as followings: red (outward motion), green (inward motion) and blue (static).

**d,** The dihedral angle between the TMD and CHD interfaces in the AF-structure and the cryo-EM structure are 83° and 86.5°, respectively. The 3.5° rotation between these two states suggest that domain rearrangement occurs during conformational conversion. We speculate that the domain re-organization is coupled to channel gating.

**e,** Malate promote the channel conversion from the basal to the activated state.

#### Extended Data Table 1

#### PSI-BLAST analysis of QUAC/ALMT-related proteins[a]

| Family /Subfamily | Representative expansion sequence | GenBank ID | -log($E_{cut}$) | Sequence |
|---|---|---|---|---|
| QF1 | *Arabidopsis thaliana* ALMT9 | NP_188473.1 | 160 | 572 |
| QF2 | *Arabidopsis thaliana* ALMT2 | NP_172320.1 | 160 | 1171 |
| Others[b] | | | | 13 |
| Total | | | | 1756 |
| QF1A | *Arabidopsis thaliana* ALMT9 | NP_188473.1 | 180 | 466 |
| QF1B | *Oryza sativa Japonica* ALMT9 | XP_015618041.1 | 180 | 106 |
| Sub-total | | | | 572 |
| QF2A | *Arabidopsis thaliana* ALMT2 | NP_172320.1 | 180 | 805 |
| QF2B[c] | *Arabidopsis thaliana* QUAC1 | NP_193531.1 | 180 | 244 |
| QF2C | *Helianthus annuus* ALMT4 | XP_022041099.1 | 180 | 122 |
| Sub-total | | | | 1171 |

[a] Procedures for the PSI-BLAST analysis of the QUAC/ALMT superfamily:

- NCBI PSI-BLAST: Thirteen expansion sequences used to define the QUAC/ALMT superfamily were as following: *At*QUAC1/ALMT12 (NP_193531.1), *At*ALMT1 (NP_172319.1), *At*ALMT2 (NP_172320.1), *At*ALMT3 (NP_173278.1), *At*ALMT4 (NP_173919.1), *At*ALMT5( NP_564935.1), *At*ALMT6 (NP_179338.1), *At*ALMT7 (NP_001324626.1), *At*ALMT8 (NP_187774.1), *At*ALMT9 (NP_188473.1), *At*ALMT10 (NP_001319836.1), *At*ALMT13 (NP_199472.1) and *At*ALMT14 (NP_199473.1). The *At*ALMT11 was not included, as it has only 152 amino acid in length, containing less than 6 TM and lacking the putative C-terminal helical domain.

The expansion sequences were run against the NCBI Protein Reference Sequence (refseq_protein), in which the green plants (taxid:33090) and an E-value $\leq 5\times10^{-3}$ were selected. Each sequence reached convergence at the 4$^{th}$ or 5$^{th}$ PSI-BLAST run, having generated ~2000 sequences. The resulted thirteen pools were merged to obtain a non-redundant set (2,061 sequences), to which some further purification was applied: i) removal of sequences not having a starting Met or containing non-amino acid X; ii) removal of sequences containing keyword "partial" in the sequence annotation; iii) removal of sequences <400 residues, unless 6 transmembrane segments confirmed by TMHMM 2.0. This resulted in a set of 1,843 sequences, which were subjected to the prediction of TM segments using THHMM 2.0. The sequences with fewer than 5 apparent TM helices were checked by *Gm*GUAC1 structure-based multiple sequence alignment with CLUSTAL_W, and sequences obviously lacking some TM helices, or having deletion/insertion in the middle of TM or other helix in the C-terminal domain were removed. This reduced to a final pool of 1,756 sequences for family/subfamily classification.

➢ Local PSI-BLAST:

The 1,756 identified sequences were used to establish a local PSI-BLAST database into which various sequences were expanded at different $E_{cut}$ levels. We found that at $E_{cut} = 10^{-160,}$ there were two distinct families plus several others. And subfamilies were divided at the level of $E_{cut} = 10^{-180}$.

[b]The other leftover sequences include 9 from *Physcomitrium patens*, 3 from *Selaginella moellendorffii*, and 1 from *Chlorella variabilis*.

[c]In QF2B, after removing redundant sequence > 95% sequence identity, a set of 155 sequences was achieved. We further checked the sequences by *Gm*GUAC1 structure-based multiple sequence alignment with CLUSTAL_W, and sequences obviously lacking some TM helix, or having deletion/insertion in the TM or helices in the C-terminal domain were removed. A final set of 137 sequences was achieved and used for sequence conservation analysis in Figure 2C.

**Extended Data Table 2**
**Statistics of data colletion, image processing and model building**

| | |
|---|---|
| Sample | *Gm*QUAC1 |
| PDB | |
| EMDB | |
| Data Collection | |
| Microscope | Titan Krios G2 |
| Voltage [kV] | 300 |
| Dectector | Gatan K2 Summit |
| Energy filter width [eV] | 20 |
| Automation software | SerialEM |
| Pixel size [Å/pixel] | 1.04 |
| Electron dose [$e^-$/Å$^2$] | 60 |
| No. of frames | 32 |
| Defocus range [µm] | -1.2 ~ -2.2 |
| Reconstruction | |
| Software | cryoSPARC 3.1 & Relion 3.0 |
| No. of particles | 169,576 |
| Symmetry | C2 |
| Map sharpening B-factor[Å$^2$] | 189 |
| Final resolution [Å] | 3.5 |
| Model Building & Refinement | |
| Building Software | Coot |
| Refinement Software | PHENIX |
| Rmsd (bond) [Å] | 0.005 |
| Rmsd (angle) [°] | 0.692 |
| MolProbity score | 2.36 |
| Model Compesition | |
| No. of residues | 820 |
| Ligands | None |
| Validation | |
| Ramachandran plot[%] | |
| Outliers | 0 |
| Allowed | 9.5 |
| Favored | 90.5 |
| Rotamer Outliers [%] | 2.01 |
